# Joint Trajectory Inference for Single-cell Genomics Using Deep Learning with a Mixture Prior

**DOI:** 10.1101/2020.12.26.424452

**Authors:** Jin-Hong Du, Tianyu Chen, Ming Gao, Jingshu Wang

**Author notes:** Equal contribution (Jin-Hong Du, Tianyu Chen).

## Abstract

Trajectory inference methods are essential for analyzing the developmental paths of cells in single-cell sequencing datasets. It provides insights into cellular differentiation, transitions, and lineage hierarchies, helping unravel the dynamic processes underlying development and disease progression. However, many existing tools lack a coherent statistical model and reliable uncertainty quantification, limiting their utility and robustness. In this paper, we introduce VITAE (**V**ariational **I**nference for **T**rajectory by **A**uto**E**ncoder), a novel statistical approach that integrates a latent hierarchical mixture model with variational autoencoders to infer trajectories. The statistical hierarchical model enhances the interpretability of our framework, while the posterior approximations generated by our variational autoencoder ensure computational efficiency and provide uncertainty quantification of cell projections along trajectories. Specifically, VITAE enables simultaneous trajectory inference and data integration, improving the accuracy of learning a joint trajectory structure in the presence of biological and technical heterogeneity across datasets. We show that VITAE outperforms other state-of-the-art trajectory inference methods on both real and synthetic data under various trajectory topologies. Furthermore, we apply VITAE to jointly analyze three distinct single-cell RNA sequencing datasets of the mouse neocortex, unveiling comprehensive developmental lineages of projection neurons. VITAE effectively reduces batch effects within and across datasets and uncovers finer structures that might be overlooked in individual datasets. Additionally, we showcase VITAE’s efficacy in integrative analyses of multi-omic datasets with continuous cell population structures.

## Introduction

Single-cell genomics has emerged as an indispensable tool for biologists seeking to unravel cellular diversities and understand cell activities [1]. Many biological processes, including differentiation, immune response, and cancer progression, can be comprehended as continuous dynamic changes within the space of cell types or cell states [2]. Instead of being limited to discrete cell types, cells often display a continuous spectrum of states and actively transit between different cellular states. To tackle this complexity, trajectory inference (TI) has been developed as a computational approach to explore the sequential states of measured cells using single-cell genomics data [3].

While various computational tools have been developed for TI [4–7], there remains a lack of clear definition for the terms “trajectory” and “pseudotime” that can be identified and estimated from single-cell genomics data. Single-cell sequencing data only captures static snapshots of single-cell omics at specific time points, which may make it challenging to uncover the true temporal lineages sought after by biologists [8, 9]. As a consequence, it is beneficial to construct an explicit and coherent statistical model on single-cell sequencing data that incorporates the trajectory structure. Such a model would also allow for discussions about the efficiency and accuracy of TI methods. Moreover, with the expansion of single-cell sequencing data [10], multiple datasets from different individuals or labs, measuring different omics are increasingly available for the same or closely related biological processes. However, to the best of our knowledge, no existing TI method can simultaneously integrate multiple datasets and infer a jointed underlying trajectory while preserving true biological differences across datasets.

In this paper, we present a novel method **VITAE** (**V**ariational **I**nference for **T**rajectory by **A**uto**E**ncoder) for performing TI. VITAE combines a graph-based hierarchical mixture model, which incorporates the trajectory structure within a low-dimensional latent space, with deep neural networks that non-linearly map the latent space to the high-dimensional observed data. We offer a clear definition of both the trajectory backbone and the pseudotime while also providing scalable uncertainty quantification of the estimates through approximate Bayesian inference. To train VITAE, we employ a modified variational autoencoder and introduce the Jacobian regularizer, an additional penalty in the loss function that significantly enhances the stability and accuracy of our algorithm. Moreover, VITAE can reliably perform joint trajectory analysis of multiple datasets by flexible modeling and effectively accounting for confounding variables such as batch effects. With VITAE, we provide comprehensive joint trajectory analyses of three single-cell RNA sequencing (scRNA-seq) datasets on mouse neocortex. We also performed a joint analysis integrating scRNA-seq with the single-cell assay for transposase-accessible chromatin by sequencing (scATAC-seq) of hematopoietic stem cells.

## Results

### Method overview

We begin our approach by modeling the trajectory backbone of cells as a subgraph of a complete graph 𝒢 = (𝒩, ℰ), where each of the *k* vertices represents a distinct cell state, and each edge represents a potential transition between two states. As shown in Fig. 1a, to capture the scenario where a cell *i* can either belong to a specific state or be in the process of transitioning between states, we define its position 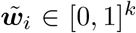 on the graph as either on a vertex if it is in a steady state or on an edge if it is undergoing a transition. Specifically, we define

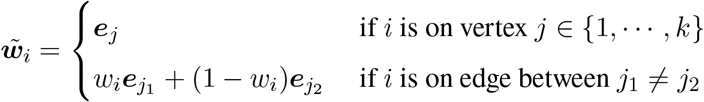

where ***e***_*j*_ is a one-hot vector with the *j*-th element as 1 and all other elements as 0. The value *w*_*i*_ ∈ [0, 1] represents the relative position of cell *i* along an edge, indicating its progress along the transition. The goal of trajectory inference is to identify the trajectory backbone, which is a subgraph containing cells positioned on both edges and vertices, as well as to estimate the cell positions 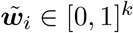, representing the pseudotime of cells along the underlying developmental process (see Methods). By quantitatively defining the developmental trajectory and pseudotime as features of a graph, VITAE offers enhanced interpretability and flexibility compared to other existing methods.

**Figure 1:**
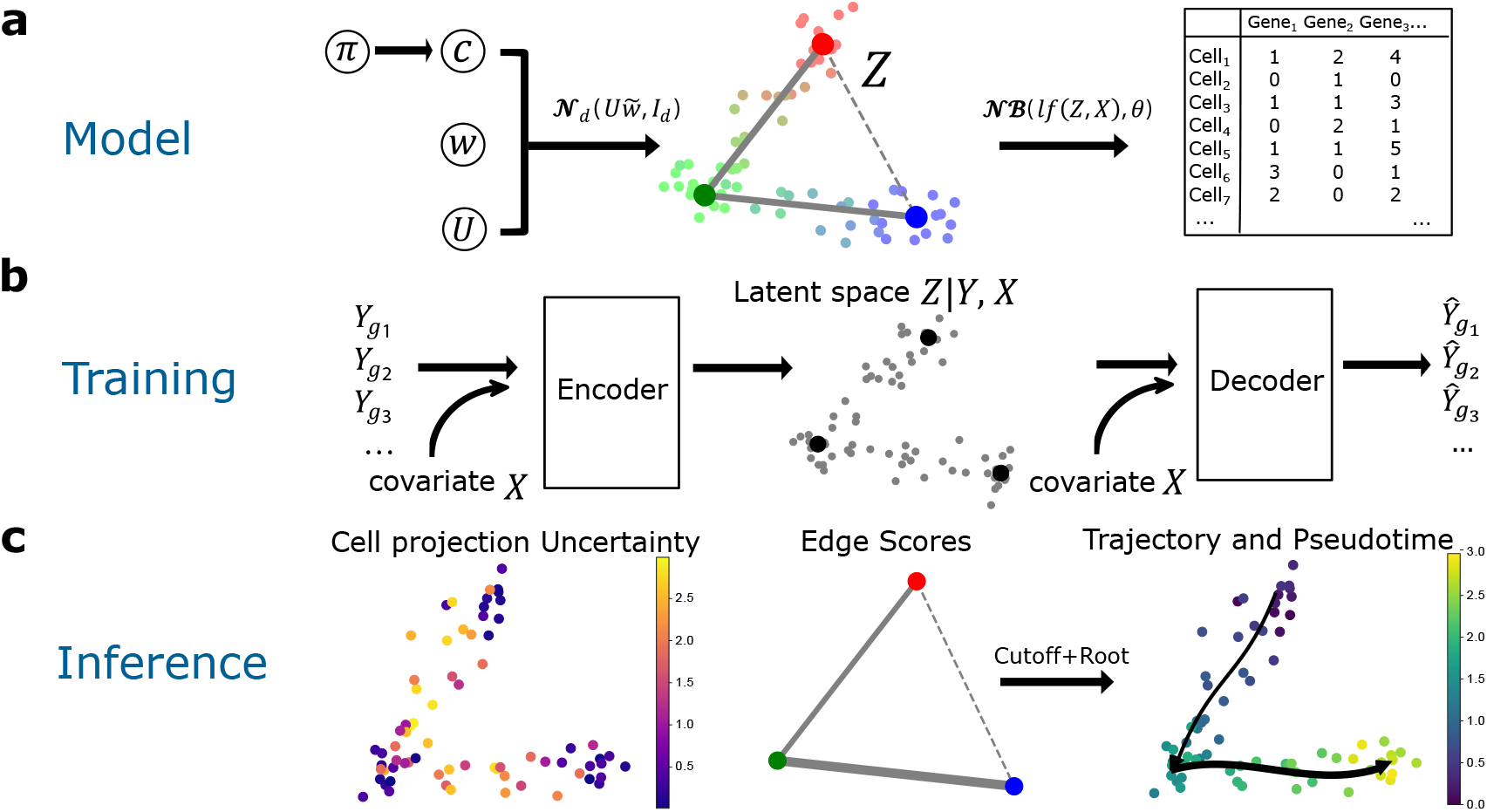
An overview of VITAE. **(a)** Model framework: A hierarchical model is used on the latent representations of cells, which are subsequently nonlinearly mapped to the high-dimensional observed count data. **(b)** Model training: Neural networks are used to approximate the nonlinear mapping to the observed data (decoder) as well as the posterior distributions of the latent space (encoder). This process also allows for adjustment for confounding covariates. **(c)** Trajectory inference: By utilizing posterior approximations, VITAE infers the trajectory backbone by assigning edge scores (represented as the width of gray lines), quantifies the uncertainty in estimated cell positions along the trajectory, and derives estimates for the pseudotime of each cell.

To connect the observed single-cell sequencing data with the underlying trajectory backbone graph in the latent space, we model the observed count *Y*_*ig*_ for any cell *i* and gene *g* as negative binomial distribution:

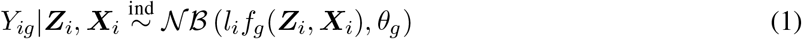

Here, *l*_*i*_ represents the library size of cell *i*, and *θ*_*g*_ is the dispersion parameter of gene *g*. The variable ***Z***_*i*_ ∈ ℝ^*d*^ represents the cell *i* in a latent lower-dimensional space, and its values are linearly dependent on the cell’s relative position along the trajectory backbone:

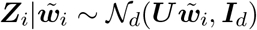

where ***U*** ∈ ℝ^*d*×*k*^ denotes the positions of the vertices on the latent space. To integrate multiple datasets, VITAE accounts for the effect of confounding variable ***X***_*i*_, which could include both discrete and continuous variables such as the dataset ID, batch indices, and cell cycle scores. We incorporate a flexible function *f*_*g*_(***Z***_*i*_, ***X***_*i*_), where *f*_*g*_(·) is represented by a neural network. By employing this setup, VITAE is capable of jointly performing data integration and trajectory inference.

VITAE is trained using a modified variational autoencoder (VAE, Fig. 1b). Based on the training dataset 𝒟_*N*_ = {(***X***_1_, ***Y***_1_), …, (***X***_*N*_, ***Y***_*N*_)}, one key idea is the incorporation of four penalty terms in the likelihood-based loss function, defined as follows:

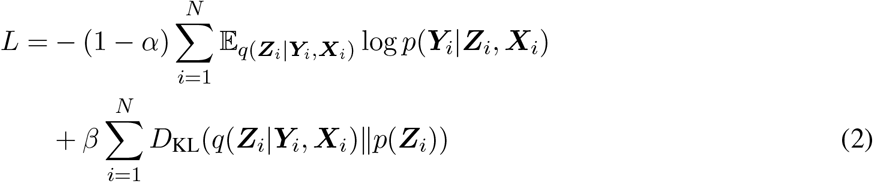

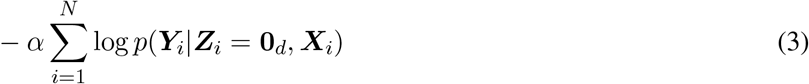

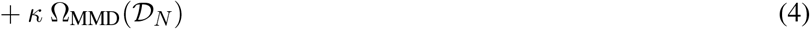

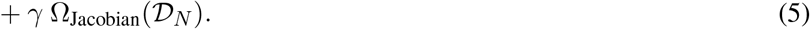

Compared to standard VAE, the prior *p*(***Z***_*i*_) contains the trajectory structure, and the term [2] that encourages the posterior positions of ***Z***_*i*_ to lie along the trajectory backbone. Then both terms [3] and [4] help adjust for the confounding variables ***X***_*i*_ and correct for batch effects. The soft penalty term [3] can effectively preserve biologically meaningful differences across datasets [11] and can also adjust for continuous confounding variables in ***X***_*i*_ such as the cell cycle scores. On the other hand, the MMD loss [4] acts as a stronger penalty, ensuring the removal of unwanted variations, and is used when cell populations across batches (e.g., replicates) are known to be exactly the same [12]. Finally, the introduction of the Jacobian regularizer [5] enhances stability in optimization [13, 14] and is a novel contribution to the analysis of single-cell sequencing data, making the estimation procedure robust to small perturbations in the observed counts. More details on the model architecture, the loss function, training and inference procedures, differential gene expression analysis, and empirical evaluation of the penalty terms are described in Methods and Appendix.

### Systematic benchmarking with real and synthetic Datasets

We systematically evaluate the performance of VITAE on 25 real and synthetic single-cell RNA sequencing datasets (Table 1), with various trajectory topologies (Fig. S.1). We examine two versions of VITAE: the original likelihood-based VITAE and its accelerated Gaussian counterpart, VITAE_Gauss (see Methods), that make VITAE scalable to handle large datasets. Our assessment involves a comparative analysis against three established TI methods: PAGA [7], Monocle 3 [15], and Slingshot [6]. To ensure an equitable evaluation, we largely adhere to the evaluation framework presented in a third-party benchmarking study [4], and the other three methods are executed through dyno [4], a standardized interface for benchmarking TI methods. Drawing inspiration from the benchmarking study [4], we employ five metrics for assessment. The GED score and IM score measure the accuracy of trajectory backbone recovery. Meanwhile, the adjusted rand index (ARI) and the generalized rand index (GRI) evaluate the precision of predicted cell positions along the trajectory, the latter being independent of clustering. Lastly, the PDT score quantifies the similarity between reference and estimated pseudotime values for cells. All five scores are normalized to a scale of 0 to 1, with higher scores indicating superior performance.

The evaluation results are summarized in Fig. 2. Both VITAE and VITAE_Gauss consistently outperform the other methods. In particular, VITAE demonstrates enhanced accuracy in recovering trajectory backbones, subsequently leading to improved pseudotime estimations. Regarding ARI and GRI, VITAE displays consistent enhancement across all synthetic datasets, although the improvements in certain real datasets are subtle due to the limited resolution of reference cell positions in real datasets.

**Figure 2:**
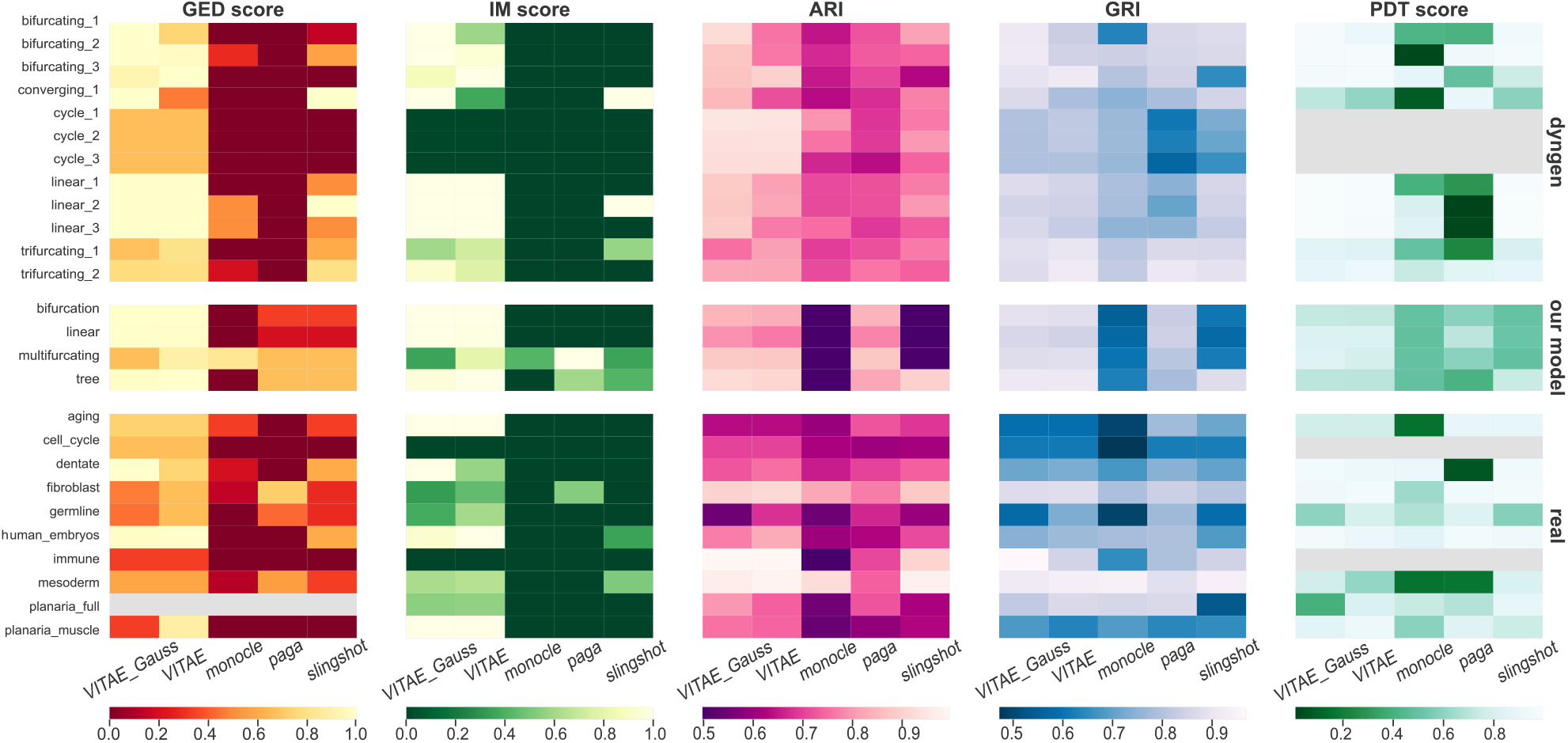
Evaluation results of VITAE and VITAE_Gauss compared with three other TI methods: Monocle 3, PAGA, and Slingshot, on 25 datasets using 5 evaluation scores. A larger score (lighter color) indicates better performance. Each row represents one dataset (Table 1). The PDT scores for datasets with the *cycle* and *disconnected* trajectory topology are not computed as the pseudotime is ill-defined. The GED scores for the *planaria_full* dataset are also not computed due to computational infeasibility. The scores for our methods are averaged over 100 runs with different random seeds.

We also assess the effectiveness of our approach in differential gene expression analysis and compare it with two alternative approaches, tradeSeq [16] and PseudotimeDE [17]. The evaluation is conducted using synthetic data generated by dyntoy from the dynverse toolbox [18] (Appendix S.3). Our approach achieves better control of false discovery rates, while maintaining comparable power and exhibiting enhanced computational efficiency when compared to the alternative methods. (Fig. S.2).

### Joint analyses of two single-cell RNA sequencing data on developing mouse neocortex

The six-layered neocortex forms the physical center for the highest cognitive functions in mammals [19]. It has been shown that the neuroepithelial cells (NEC) transition into the radial glial cells (RGC), while cortical projection neuron types are generated sequentially by RGCs and intermediate progenitor cells (IPC) [20, 21]. In our study, we employ VITAE to infer a unified trajectory framework through the integration of two distinct single-cell RNA sequencing datasets pertaining to the developing mouse neocortex. Specifically, we delve into Yuzwa’s dataset [22], encompassing 6, 390 cortical cells sampled from mouse embryos at E11.5, E13.5, E15.5, and E17.5 time points. Additionally, we incorporate Ruan’s dataset [23], which has 10, 261 cells derived from the mouse embryo cortex at E10.5, E12.5, E14.5, E15.5, E16.5, and E18.5 (Table 2). Thus, these datasets capture cells from disparate developmental days within the same brain region. By jointly analyzing these datasets, we unlock the potential to investigate the dynamic progression spanning a comprehensive developmental timeline, from E10.5 to E18.5 in mice.

To evaluate the performance of VITAE, we conducted a comparison between VITAE and an alternative approach, starting with integration using Seurat CCA [24], followed by trajectory inference using Slingshot on the integrated embeddings. While both VITAE and Seurat CCA are effective at cell mixing and demonstrate comparable retention of reference cell types post-integration, VITAE excels in providing a more refined trajectory inference on the shared structure (Fig. 3ab). In particular, VITAE successfully identified the primary cell lineage, denoted as *NEC-RGC-IPC-Neurons*, while excluding microglia cells, pericytes, Pia, and interneurons from the Ruan dataset that do not align with the main trajectory. On the contrary, Slingshot yields disorderly and ambiguous outcomes, attributed in part to subtle imperfections within the Seurat integration process and its inability to recognize disconnected cell types. To assess the stability of VITAE, we further conducted 10 repeated trials of VITAE with random initialization and observed that the inclusion of the Jacobian regularizer substantially reduces the variability in our estimated hierarchical model of the trajectory backbone (Fig. S.3).

**Figure 3:**
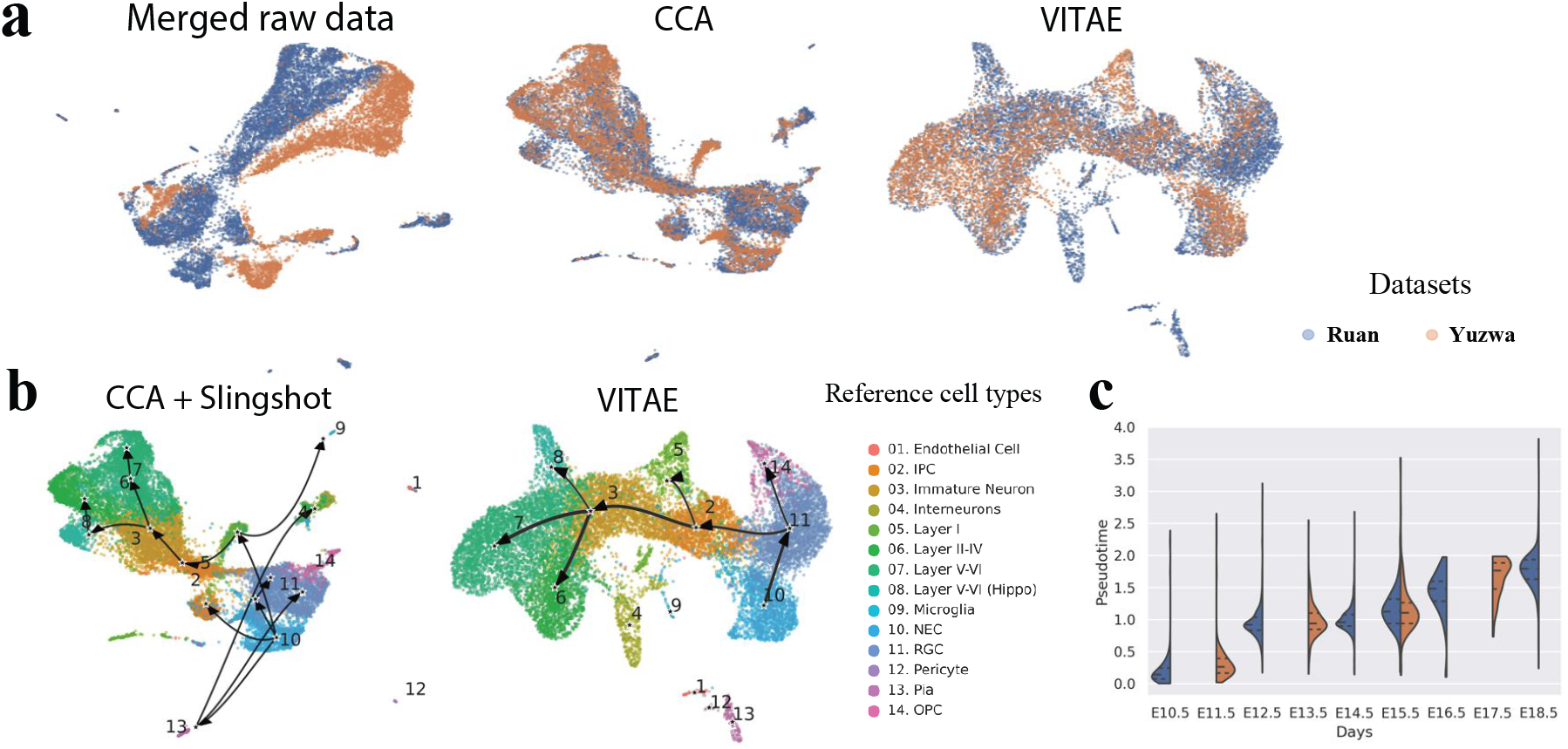
UMAP visualizations of the mouse neocortex in Yuzwa and Ruan datasets. **(a)** Low-dimensional embeddings colored by dataset sources. We compare the UMAP plots of the cells from the merged raw data, after Seurat CCA integration and by VITAE. The cell cycle scores are adjusted for both Seurat CCA and VITAE. **(b)** The estimated trajectories from Seurat CCA + Slingshot (left) compared with that from the accelerated Gaussian version of VITAE (right). Cells are colored by their reference cell types given in the original papers. For VITAE, the width of each edge is proportional to its edge score. **(c)** The violin plots of estimated pseudotime by VITAE across different embryonic days in the sub-trajectory *NEC-RGC-OPC*.

Regarding uncertainty quantification, VITAE provides edge scores for each inferred trajectory edge (Fig. 3b, displayed as the edge widths), revealing our confidence in the existence of each edge. Additionally, VITAE estimates mean square errors for the projection of each cell onto the inferred trajectory structure (Fig. S.4a).

Moreover, we observe robust correlations between the developmental days of cell collection and the pseudotime estimation along the sub-trajectory *NEC-RGC-OPC* (Fig. 3c, Fig. S.4b-d). Although cells collected on even days are from the Ruan dataset, and those collected on odd days mainly originate from the Yuzwa dataset, VITAE effectively aligns cells along pseudotime gradually according to their developmental days. This indicates VITAE’s ability to preserve biologically meaningful distinctions across datasets while simultaneously learning a shared trajectory structure.

### Integration with the mouse cerebral cortex atlas

Extending our analysis to encompass a more intricate system, we perform an additional joint trajectory inference analysis involving the Ruan and Yuzwa datasets in conjunction with the Di Bella dataset [25] – a comprehensive mouse neocortex atlas with 91,648 cells that contains daily samples from the neocortex throughout embryonic corticogenesis and at the early stage of postnatal development. Due to differences in sequencing technologies, the observed Di Bella dataset exhibits significant differences in comparison to the Ruan and Yuzwa datasets (Fig. S.5). Additionally, there are substantial batch effects across cells gathered from various embryonic days within the Di Bella dataset, even spanning different technical replicates. We aim to use VITAE to mitigate these batch effects within the Di Bella dataset and project cells from the Ruan and Yuzwa datasets onto the Di Bella dataset, thereby facilitating a more comprehensive and integrated analysis.

We compare VITAE with the alternative approach using Seurat CCA for integration and Slingshot for trajectory inference. Additionally, we benchmark with Monocle 3 [15], as employed in the original paper [25], to undertake trajectory inference. Monocle 3 facilitates data integration as its preliminary step and we leverage this to integrate the three datasets before embarking on trajectory inference when using Monocle 3. In alignment with the outcomes presented in the preceding section, VITAE can reveal a more refined joint trajectory structure when juxtaposed against the other two methodologies (Fig. 4a), recovering the main branches identified in the original paper [25] of Di Bella dataset.

**Figure 4:**
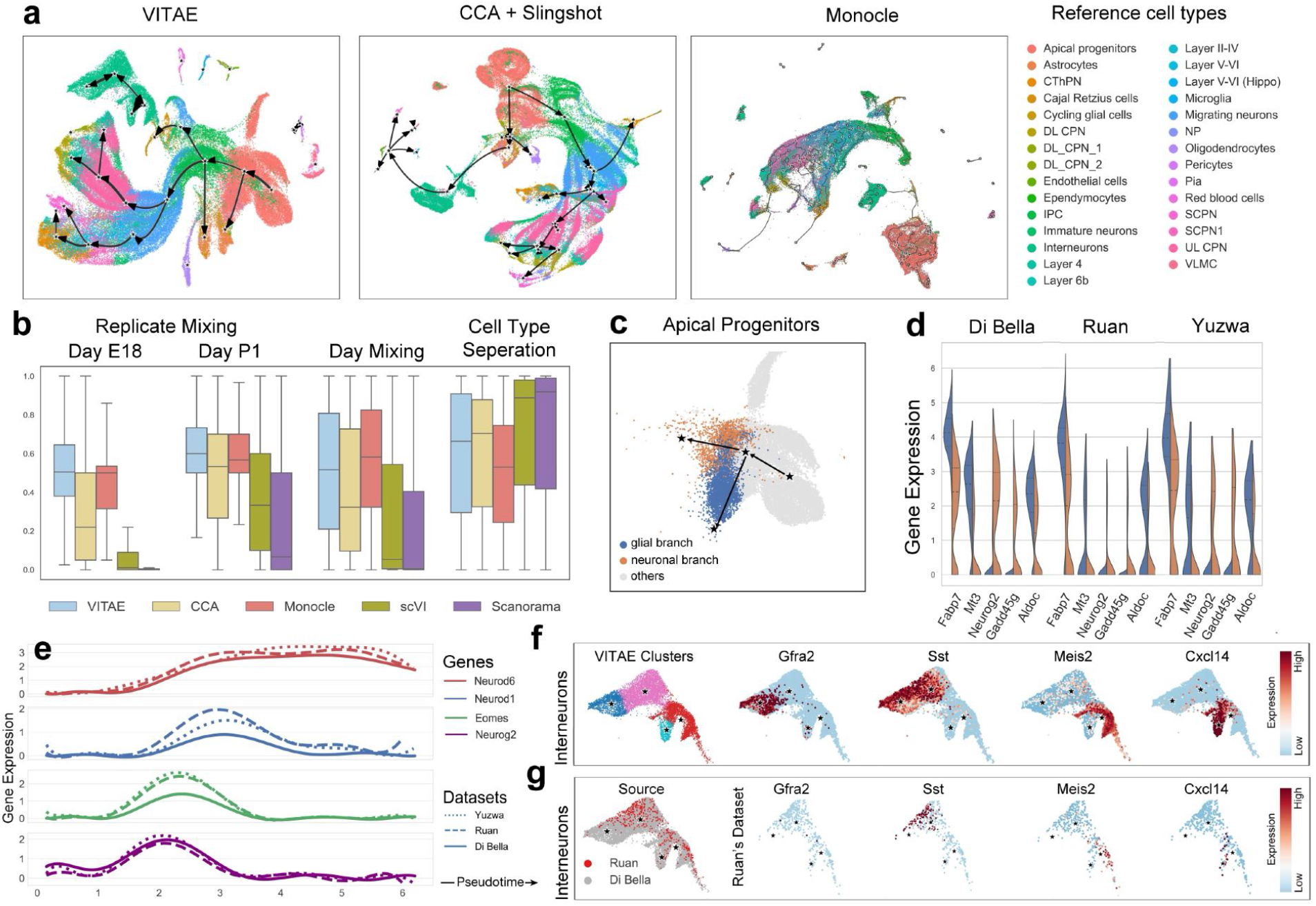
Integrative trajectory analysis on three mouse brain datasets. **(a)** The estimated trajectories on the merged dataset using different methods. Cells are colored by their reference cell types provided in the original paper. **(b)** Comparison of data integration efficacy across three methods measured by the replicate mixing scores, day mixing scores, and cell type separation scores. A higher score indicates better performance. **(c)** Using VITAE, some APs are projected onto the glial branch (blue) or the neural branch (yellow). All other APs are colored gray. **(d)** The violin plots of the selected differentially expressed gene expressions between the APs in the glial (blue) and neuronal (orange) branches. Tests are done by combining all three datasets while gene expressions are visualized for each dataset separately. **(e)** Smoothed curve for gene expression change along the estimated pseudotime. **(f)** Four interneuron subtypes identified by VITAE. VITAE automatically identifies four vertices within the interneurons though a single initial vertex is given. **(g)** Interneurons from the Ruan dataset positioned within the four interneurons subtypes.

We also assess the efficacy of data integration across the three methodologies, comparing them with two additional methods, scVI [26] and Scanorama [27], which are specifically designed for data integration. The complexity of data integration arises from the fact that cells originating from different embryonic days have inherent biological differences, but are also collected in different batches. As the cells exhibit continuous cell states, batch effects can be comparable or even stronger in magnitude to biological differences across embryonic days. Therefore, we can not simply treat embryonic days as batches and explicitly adjust for the differences. Without explicitly adjusting for differences in embryonic days, we observe that VITAE performs the best in reducing batch effects across embryonic days while retaining genuine biological variances (Fig. 4b). Considering the presence of two technical replicates for Day E18 and Day P1 within Di Bella’s dataset, VITAE has the highest median replicate mixing scores (Appendix S.4.2) in both the two replicates. In addition, Monocle 3 exhibits the poorest performance in terms of cell type separation, implying an excessive removal of biologically meaningful cell-type-specific signals. While Seurat CCA, scVI, and Scanorama achieve better or comparable cell type separation than VITAE, they inadvertently maintain a complete separation between cells from distinct embryonic days, indicating excessive retention of batch effects (Figs. S.5 and S.6).

Leveraging the inferred trajectory generated by VITAE, we analyze branching within the apical progenitors (APs), comparing the APs projected onto the glial branch and those projected onto the neuronal branch (Fig. 4c). Through differential analysis, we identify enrichment of radial glia markers (*Fabp7, Mt3, Dbi, Aldoc, Ptprz1*) within the APs situated along the glial branch (Fig. 4d, Fig. S.7a). Conversely, genes such as *Neurog2* and *Hes6* have higher expression in APs projected onto the neural branch, suggesting a potential state of primed neurogenesis. These findings closely mirror the outcomes of the analysis conducted in the original Di Bella dataset paper. However, the convenience of our integrated VITAE framework significantly facilitates the execution of this analysis. Additionally, we are able to confirm that this divergence trend within the AP population holds consistent across all three datasets.

We also visualize the expressions of IPC marker *Eomes*, neurogenesis marker *Neurog2*, migration-associated gene *Neurod1*, and neuronal differentiation gene *Neurod6* along the estimated pseudotime (Fig. 4e). We observe that the expression of *Neurog2* precedes that of *Neurod1* and *Neurod6*, confirming its role as a proneural gene within the dorsal telencephalon [23]. Despite the presence of lab-specific batch effects, these genes exhibit coherent patterns across all three datasets.

Finally, we focus on subtype analysis of the interneurons. With the joint analyses of all three datasets, VITAE can automatically identify four biologically different interneuron cell subtypes (Fig. 4f, Fig. S.7b). What’s particularly noteworthy is that while three of these subtypes align with those previously identified through clustering in the original Di Bella paper that analyzes solely its own atlas data, VITAE reveals an additional subtype characterized by the expression of *Gfra2*. This subtype potentially represents the GABAergic interneuron population, as suggested by previous studies [28, 29]. Furthermore, VITAE reliably positions the small population of interneurons from the Ruan dataset within these four identified subtypes (Fig. 4g). Only a minority of interneurons from the Ruan dataset exhibit characteristics associated with the *Gfra2+, Sst+*, and *Meiz2+* subtypes, as they predominantly emerge during the later stages of embryonic days (Fig. S.7c). Without the joint analysis facilitated by VITAE, it would be challenging to identify these interneurons solely from the Ruan dataset.

### Differential analysis along estimated trajectories

VITAE is also designed to identify differentially expressed genes along any estimated sub-trajectory. We focus on examining gene expression dynamics along the sub-trajectory spanning RGC to IPC to migrating neurons (Fig. 5a). Despite that cells are from three distinct sources for this analysis, a consistent pattern emerges in the phase transitions of these differentially expressed genes across all three datasets (Fig. 5b). This confirms VITAE’s ability to align cells originating from disparate datasets while preserving their inherent biological information.

**Figure 5:**
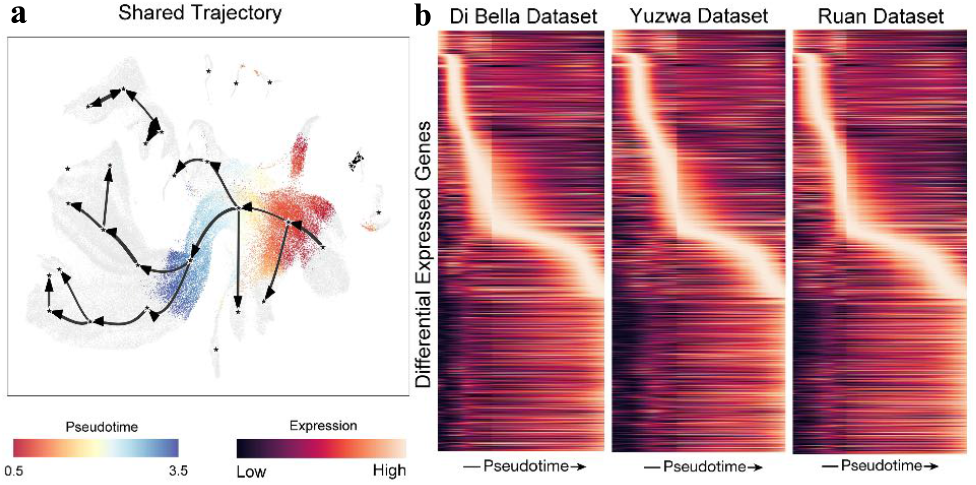
Testing for differentially expressed genes along a sub-trajectory on three mouse brain datasets. **(a)** Highlight of the sub-trajectory containing the apical progenitors, migrating neurons, and immature neurons with cells colored by pseudotime. **(b)** The heatmaps of DE gene expressions for the sub-trajectory along the inferred pseudotime ordering of the cells in each dataset. The DE genes are identified by performing quadratic regression of the gene expressions on the standardized pseudotime ordering with covariates adjusted (*t*-test, adjusted *p*-value*<* 0.05).

### Joint trajectory analysis on scATAC-seq and scRNA-seq datasets

We further apply VITAE to conduct a combined trajectory analysis of cells using both scRNA-seq and scATAC-seq data obtained from the same human hematopoiesis population of healthy donors [30]. The joint analysis of the two modalities can help to identify the cell types within the ATAC-seq data and infer active regulators for the RNA-seq data without requiring the two modalities to be measured on the same individual cells [31]. We use VITAE to simultaneously integrate the two modalities and infer a developmental trajectory for hematopoietic stem cells (HSC). VITAE successfully achieved a uniform mixing of cells from both modalities while preserving the distinction between different reference cell types (Fig. 6a, Fig. S.8a). VITAE reveals the two major lineages (Fig. 6a), including the B cell lineage (*HSC-CMP*.*LMPP-CLP-Pre*.*B-B*) and the monocytic lineage (*HSC-CMP*.*LMPP-GMP-CD14*.*Mono*.*1-CD14*.*Mono*.*2*). We observed consistent patterns of continuous cell type transitions along the pseudotime in both modalities and within these two lineages (Fig. 6b, Fig. S.8b). We also compare with an alternative approach VIA [32] and observe the superior performance of VITAE in learning a clean and reliable shared trajectory structure across modalities (Fig. S.8c).

**Figure 6:**
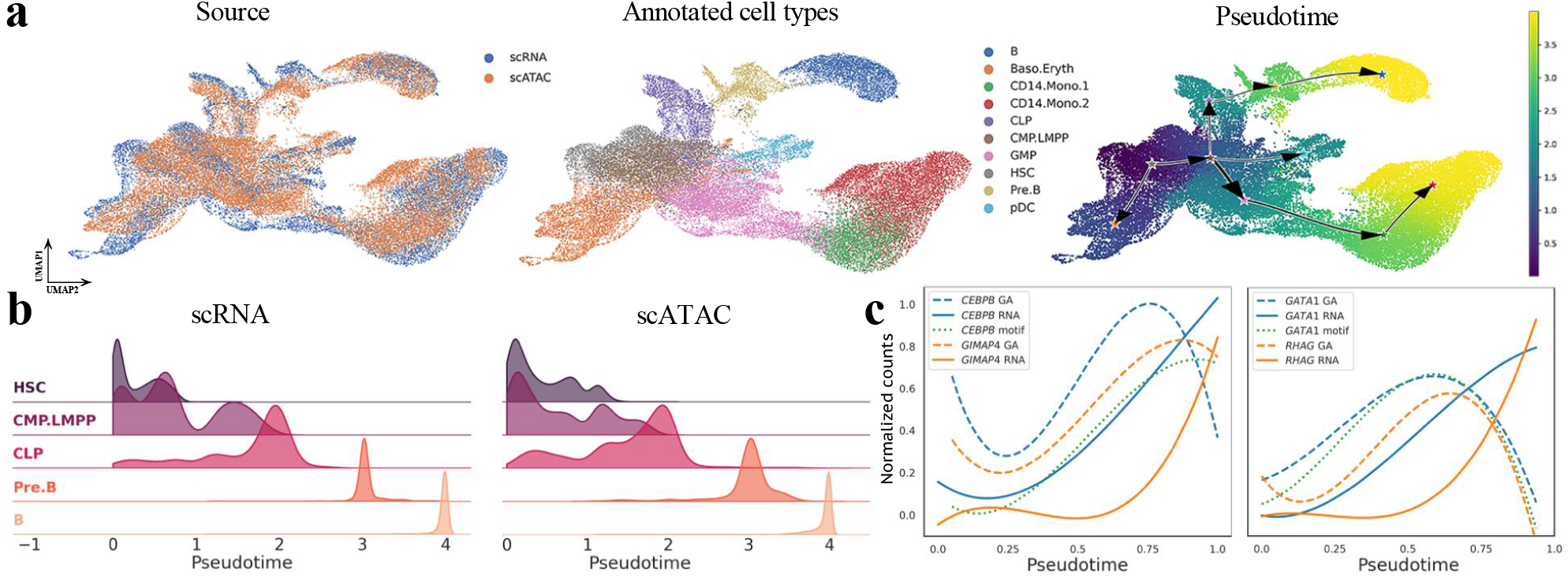
Integrative trajectory inference of multi-omic human hematopoiesis data. **(a)** UMAP visualization of VITAE’s low-dimensional embedding of cells, colored by source, annotated cell types, and pseudotime. **(b)** Distribution of cells per cell type along the estimated pseudotime of the B cell lineage. **(c)** Lineage dynamics of the *CEBPB* and *GATA1* genes. The gene activity (GA) scores, TF (motif) scores, and gene expressions precede the gene expressions of their target genes, *GIMP4* and *RHAG* respectively, along the estimated pseudotime in each lineage.

By joint profiling of chromatin accessibility and gene expression within the same individual cells, earlier papers suggest that changes in chromatin accessibility may prime cells for lineage commitment [33]. By projecting cells onto the shared trajectory, VITAE effectively aligned cells with only one of the two omics, allowing us to visualize similar regulatory dynamics. For instance, we examined the gene *CEBPB*, a key transcriptional factor (TF) in monocytic development, and observed a clear progression of its gene activity score, gene expression level, and TF activity score, followed by the expression level of its target gene *GIMP4*. Similar trends were also identified for the gene *GATA1*, a pivotal TF in the erythroid lineage, and its target gene *RHAG*. These findings underscore the power of VITAE in elucidating regulatory relationships within multi-omics data.

## Discussion

In this paper, we propose VITAE, a hierarchical mixture model combined with VAE, for trajectory inference. Compared with existing TI methods, VITAE provides a coherent probabilistic framework to explicitly define estimable trajectories and pseudotime while accommodating diverse trajectory topologies and effectively accounting for confounding covariates. VITAE harnesses the power of variational autoencoders to approximate posterior quantities, facilitating the inference of trajectory backbones and cell positions from these approximations. Moreover, VITAE incorporates a Jacobian regularizer as a penalty to enhance the model’s estimation robustness.

While numerous tools exist for integrating multiple scRNA-seq datasets, they often struggle when dealing with cells exhibiting continuous trajectory structures that do not neatly fit into distinct cell types. Our approach, which seamlessly combines trajectory analysis with data integration, excels at regularizing the latent space, resulting in a cleaner joint trajectory structure while preserving biologically meaningful distinctions in each dataset.

However, it’s essential to acknowledge the limitations of our approach. First, the approximate variational inference employed to infer trajectories and quantify cell position uncertainties overlooks the estimation uncertainties inherent in our encoder and decoder. Consequently, while our uncertainty quantification offers valuable insights, it may not precisely reflect the true posterior uncertainties. Additionally, though VITAE learns a joint trajectory structure that can contain both shared and dataset-specific sub-trajectories, there is no test to distinguish between the two. As a potential solution, we can integrate the estimated pseudotime and trajectory from VITAE with Lamian [34] or condiments [35], which are specifically designed to identify condition-specific sub-trajectories. Finally, our current framework exclusively accommodates multi-omics data with identical input dimensions. Future work will focus on extending VITAE to effectively leverage multi-omics data with distinct features for joint trajectory inference.

## Materials and Methods

### Definition of the trajectory backbone and pseudotime

In VITAE, the trajectory backbone, denoted as ℬ, is defined as a subgraph of 𝒢 that only includes edges with positive proportions of cells:

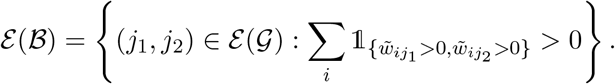

To determine the pseudotime of a cell, we first define the pseudotime for each vertex. Given a root vertex *k*_0_, we first generate a directed trajectory backbone 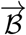 (Appendix S.1). Let each edge 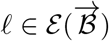 be associated with a duration *b*_*ℓ*_ (with a default value of *b*_*ℓ*_ = 1 for all edges). The pseudotime for vertex *j* is defined as:

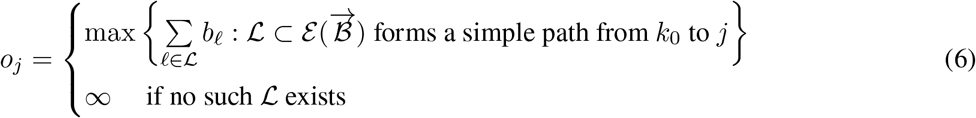

where a simple path is a path that does not have any repeating vertices. Let ***o*** = (*o*_1_, · · ·, *o*_*k*_), then for a specific cell *i* with its position 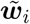 on the graph, its pseudotime is defined as:

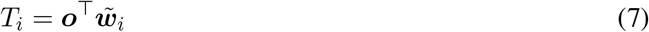

Intuitively, *T*_*i*_ equals the pseudotime of the vertex the cell is on or the weighted average of the pseudotime of the two vertices of the edge to which the cell belongs.

By our definition, any vertex that precedes another vertex on a simple path that starts from *k*_0_ has a smaller pseudotime than the other vertex. Thus our definition guarantees a meaningful ordering of the cells when the directed trajectory backbone 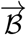 does not contain any cycles. If 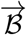 contains cycles, the pseudotime of the cells remains well-defined under our definition but may not be biologically meaningful.

### Hierarchical model for the distributions of gene expressions

In equation (1), we model the observed counts, such as those from scRNA-seq with Unique Molecular Identifiers (UMI), using a Negative Binomial (NB) distribution with gene-specific dispersion parameters *θ*_*g*_. This choice is suitable for capturing the stochasticity in scRNA-seq data, accounting for both biological and technical noise [36]. However, for certain cases where the NB distribution may not be adequate, such as scRNA-seq without UMI or single-cell ATAC sequencing data with numerous excessive zeros, we adopt the assumption that the observed counts follow zero-inflated Negative Binomial (ZINB) distributions, as proposed by [37]:

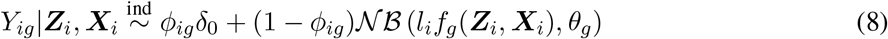

where *ϕ*_*ig*_ = *h*_*g*_(***Z***_*i*_, ***X***_*i*_) is the zero inflation probability with *h*_*g*_(·) being unknown non-linear functions.

To facilitate model estimation, we introduce a hierarchical prior on the cell positions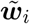. Let the latent category variable *c*_*i*_ take values from the set 1, 2, …, *K*, where *K* = *k*(*k* + 1)*/*2 represents the total count of potential edges and vertices within the graph 𝒢. Then *c*_*i*_ ∼ Multinomial(1, ***π***) serves as an indicator of the chosen edge or vertex by the cell, with ***π*** being empirically estimated during the model training step. If the cell is positioned on an edge, we assume that its relative placement along the edge has a uniform prior: *w*_*i*_ ∼ Uniform(0, 1). For more mathematical details, refer to Appendix S.1.1.

To reduce the computational cost, we also present an accelerated Gaussian version of VITAE. In this version, the observed counts of the *G* genes are replaced with the top *R* (*R* = 64 by default) principle component (PC) scores ***F***_*i*_. The distribution of ***F***_*i*_ is then approximated by Gaussian distributions. This modification allows VITAE to scale effectively and achieve a computational efficiency comparable to other trajectory inference methods.

### Penalties in the loss function

As demonstrated in the main text, VITAE’s loss function comprises four penalty terms [2]-[5]. While the marginal density function *p*(***Z***_*i*_) in [2] involves a complicated hierarchical mixture model, it possesses a tractable closed-form representation (Appendix S.2), facilitating straightforward evaluation. By default, we assign *α* = 0.1 to the soft penalty [3], and incorporate the MMD loss [4] solely in scenarios involving technical replicates or when the soft penalty [3] is insufficient and a stronger penalty is necessary.

For the definition of MMD loss, consider two cell groups 𝒫_1_ and 𝒫_2_, each with *n*_1_ and *n*_2_ cells, respectively. To eliminate all differences between the two groups, the MMD loss is characterized in the latent space by the following formulation:

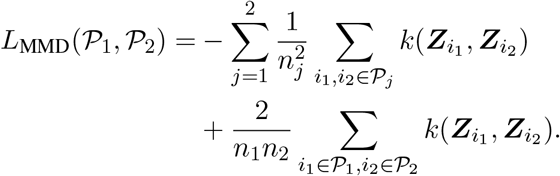

where *k*(·, ·) is the multi-scale RBF kernel proposed by [38]. Consequently, the MMD penalty [4] is outlined as the cumulative MMD loss across all such pairs of cell groups:

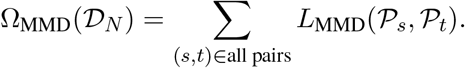

Lastly, the Jacobian regularizer [5] is defined to control the Frobenius norm of the Jacobians of latent variables ***Z***_*i*_ with respect to the input ***Y***_*i*_. Specifically, the penalty term [5] is defined as:

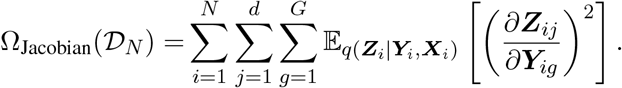

The default value for the tuning parameter *γ* is set to 1.

### Model training, estimation, and trajectory inference

We train VITAE with only highly variable genes and use normalized, log-transformed, and scaled gene expressions as input for the model. The initial parameters of VITAE are set by pretraining with *β* = 0. When cell labels are unavailable, the number of states *k* is determined by clustering the pretrained latent space (Appendix S.1.4). To estimate VITAE’s model parameters, we employ the mini-batch stochastic gradient descent algorithm, ensuring memory usage remains independent of cell count.

Using the estimated model parameters, we approximate posterior distributions for *w*_*i*_ and *c*_*i*_ by sampling from the estimated encoder output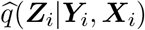. Leveraging these posterior approximations, we infer the trajectory backbone 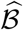 under the assumption that ℬ predominantly comprises sparse edges. Specifically, let 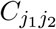 denote the edge connecting states *j*_1_ and *j*_2_. Its edge score is defined as follows:

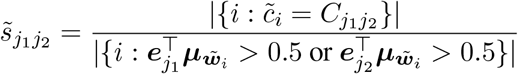

where 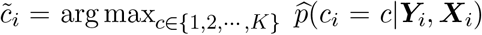 and 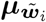 is the approximate posterior mean of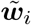. An edge is included in 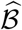 when 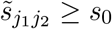 (with *s*_0_ = 0.01 as the default threshold). Subsequently, cell positions are projected onto 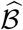 based on the aforementioned posterior approximations, which also provide uncertainty estimates for these projections.

Given the inferred trajectory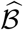, the user can manually assign a root vertex. We also provide an automatic root selection step following Tempora [39] when cells are gathered across multiple time points. With a chosen root vertex, the pseudotime of other vertices and all cells can be readily derived by substituting estimates into the definitions [6]-[7]. For a comprehensive discussion of our trajectory inference procedure from posterior approximations, refer to Appendix S.1.2.

### Differential gene expression analysis

To identify differentially expressed genes along the estimated pseudotime, we regress gene expressions of each gene on rank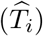, the ordering of the estimated pseudotime, via a polynomial regression, which also adjusts for confounding covariates *X*_*i*_. To account for the fact that the pseudotime 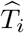 is estimated, we use a heuristic p-value calibration approach following [40]. For details, refer to Appendix S.1.3.

### Benchmarking methods

For systematic benchmarking with real and synthetic datasets, we run PAGA, Monocle 3, and Slingshot via the Dyno platform [4], which converts these TI methods’ outputs into an estimated trajectory backbone 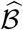, estimated cell positions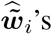, and estimated pseudotime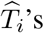. For a fair comparison, the true number of states *k*, clustering labels, and root are provided as prior information to all TI methods. For Slingshot, which requires an extra two-dimensional projection of cells as input, the UMAP [41] coordinates of cells computed following Seurat’s [24] default steps are used.

To compare our approach with tradeSeq and pseudotimeDE in differential gene expression analysis, we generate synthetic data with linear trajectories using package dyntoy from dynverse toolbox. We use Slingshot or VITAE for trajectory inference and subsequently apply our approach, tradeSeq or pseduotimeDE, to identify differentially expressed genes.

For experiments on mouse brain datasets [22, 23], we compare the trajectory inference result with Slingshot. To apply Slingshot, Seurat’s CCA is first used to integrate both datasets and regress out cell-cycle scores. Then Slingshot is applied to Seurat’s results to infer the trajectories.

For experiments on the integration of three mouse brain datasets [22, 23, 25], we compare the results with Slingshot and Monocle3. For the latter, the batch effects from the different datasets are adjusted by the function `align_cds`, and then the trajectories are learned afterward. We also compared scVI and Scanorama for data integration, providing the data sources as batch information for these methods.

For experiments on the integration of scRNA-seq and scATAC-seq human hematopoiesis data, we compare with VIA v0.1.96. Following the procedure suggested in the original paper, to apply VIA, Seurat’s CCA is first used to integrate both datasets and remove the batch effects. For details about all the benchmarking methods, refer to Appendices S.3 and S.4.

### Data, Materials, and Software Availability

- Developing mouse brain datasets. Yuzwa dataset [22] and Ruan dataset [23] are available from the GEO database under accession codes GSE107122 and GSE161690, respectively. For Yuzwa’s dataset, we only use cortically derived cells selected by the original paper. We keep the genes that are measured in both datasets (14,707 genes) and merge all 16,651 cells (6,390 and 10,261 cells for each). The cell cycle scores (`S.Score` and `G2M.Score`) of the two datasets are calculated separately by Seurat v3. As cell type labels are not provided by the Yuzwa dataset, we perform clustering using Seurat and annotate cell types using the marker genes in the Ruan dataset. The information on the collection days of both datasets is summarized in Table 2.
- Mouse cerebral cortex atlas. Di Bella’s preprocessed dataset [25] following the procedure in [25] are available in the https://singlecell.broadinstitute.org/single_cell/study/SCP1290/molecular-logic-of-cellular The cell cycle scores (`S.Score` and `G2M.Score`), cell type annotations, and collection day information are provided in the original paper. We exclude cells labeled as ‘Doublet’ and ‘Low quality cells’ and retain 91,648 cells and 19,712 genes with 24 different cell type annotations.
- Integration of the three mouse brain datasets. The merged developing mouse brain dataset and the atlas dataset are preprocessed separately by the standard procedure (`normalize_total, log1p, highly_variable_genes, scale`) using `scanpy` v1.8.3 [42] with default parameters and exclude cells labeled as ‘Doublet’ and ‘Low quality.’ Then `mouse brain` and `atlas` datasets are merged by taking the union of highly variable genes in each dataset. Then we unified different cell type annotations that refer to the same cell type (for example, ‘Intermediate progenitors’ as ‘IPC’). Finally, there are 108,299 cells and 13,183 genes. Empirically, we found that SCPN cells before and after day E16 are always projected to two different latent space regions. So, we relabeled the SCPN cells before E16 as SCPN1 to provide a more biologically meaningful center when initializing latent space.
- Human hematopoiesis data. We obtain the gene expression, peak matrix, and cell types annotation of human hematopoiesis data of healthy donors from https://github.com/GreenleafLab/MPAL-Single-Cell-2019[30]. The TF activity scores from [30] were computed using chromVAR [43]. Following the preprocessing procedure as in [44], we excluded cells labeled as ‘Unknown’ and combined the clusters with the same cell-type annotation into one label (for example, ‘CLP.1’ and ‘CLP.2’ as ‘CLP’). We retain only cell types that are on the two developmental trajectories analyzed in [44], and the ‘cDC’ and ‘CD16.Mono’ cell types are removed because of the inadequate number of cells. We calculate the highly variable genes for each dataset by scanpy and retain the union of these genes. The raw count matrices containing the selected cells and genes are then concatenated accordingly. Finally, this results in 19,309 cells for scRNA-seq data and 22,685 cells for gene activities of the scATAC-seq data for the analysis, with 4,094 genes measured.

After filtering genes and cells, we apply the standard preprocessing procedure by scanpy v1.8.3 [42] before supplying the datasets to our model.

The Python package of VITAE is publicly available at https://github.com/jaydu1/VITAE with MIT license. Python and R scripts for reproducing all results in this paper are also provided in the same repository.

## Acknowledgments

J.W. is partly supported by the National Science Foundation under grants DMS-2113646 and DMS-2238656.

