## Supplemental materials for "Joint Trajectory Inference for Single-cell Genomics Using Deep Learning with a Mixture Prior"

### Appendix

Below, we outline the structure of the rest of the supplement. Appendix S.1 introduces the proposed trajectory inference method, while the specifications of certain algorithm details are deferred to Appendix S.2. The details of simulation datasets, evaluation metrics, and experiment procedures are presented in Appendix S.3. The details of case studies on mouse brain datasets are provided in Appendix S.4. The sensitivity of hyperparameters and computational efficiency are analyzed in Appendix S.5. Lastly, the supplementary tables and extra experimental results on real data are included in Appendix S.6 and Appendix S.7, respectively.

**S.6 Supplementary tables** **50**

**S.7 Supplementary figures** **51**

### S.1 Methods

#### S.1.1 Model

**A hierarchical mixture model for the trajectory structure** Inspired by the common trajectory model proposed in [4], we start with the trajectory backbone defined on a complete graph $\mathcal{G} = (\mathcal{N}, \mathcal{E})$ with vertices $\mathcal{N}(\mathcal{G})$ and edges $\mathcal{E}(\mathcal{G})$. Let the $k$ vertices in $\mathcal{N}(\mathcal{G})$ be the distinct cell states. We use an edge between two vertices to represent the transitioning between two states. To model the scenario that one cell either belongs to a specific state or is developing from one to another, we assume that each cell is positioned either on one vertex or an edge. Specifically, let $\tilde{\boldsymbol{w}}_i \in [0,1]^k$ be the position of cell $i$ on the backbone, we have

$$\tilde{\boldsymbol{w}}_i = \begin{cases} \boldsymbol{e}_j & \text{if cell } i \text{ is on vertex } j \in \{1, \cdots, k\} \\ w_i \boldsymbol{e}_{j_1} + (1 - w_i)\boldsymbol{e}_{j_2} & \text{if cell } i \text{ is on the edge between vertices } j_1 \text{ and } j_2 \ (j_1 \neq j_2) \end{cases}$$

where $\boldsymbol{e}_j$ is a one-hot vector with $j$th element 1 and all other elements 0, and $w_i \in [0,1]$ describes the relative position of cell $i$ if it is on an edge.

The basic goal of our trajectory inference is to infer the trajectory backbone $\mathcal{B}$ - a subgraph of $\mathcal{G}$ - whose edges have positive proportions of cells:

$$\begin{aligned} \mathcal{N}(\mathcal{B}) &= \mathcal{N}(\mathcal{G}) \\ \mathcal{E}(\mathcal{B}) &= \left\{ (j_1, j_2) \in \mathcal{E}(\mathcal{G}) : \sum_i \mathbb{1}_{\{\tilde{w}_{ij_1} > 0, \tilde{w}_{ij_2} > 0\}} > 0 \right\}. \end{aligned}$$

Meanwhile, we also aim to estimate the relative positions $\tilde{\boldsymbol{w}}_i$ of each cell on this trajectory backbone and, as in other TI methods, the pseudotime order of the cells along the trajectory. Intuitively, the pseudotime corresponds to the biological progression of a cell through the dynamic process that results in the trajectory structure [3].

Here, we give a formal definition of the pseudotime in our framework. Assume that each edge $\ell \in \mathcal{E}(\mathcal{B})$ is associated with a duration (an edge weight) $b_\ell$. Without external information, we set $b_\ell = 1$. Given a root vertex $k_0$, we first generate a directed trajectory backbone $\overrightarrow{\mathcal{B}}$, by giving each edge in $\mathcal{E}(\mathcal{B})$ a direction. We start with finding the connected components $\mathcal{A} \subset \mathcal{N}(\mathcal{B})$ containing $k_0$ on the undirected trajectory backbone $\mathcal{B}$. Any edges that do not connect to a node in $\mathcal{A}$ become a bi-directional edge in $\overrightarrow{\mathcal{B}}$. If the subgraph of $\mathcal{B}$ with nodes in $\mathcal{A}$ forms a tree, then given the root $k_0$, we turn it into a directed tree starting from $k_0$. However, if the undirected subgraph of $\mathcal{B}$ with nodes in $\mathcal{A}$ contains any cycles, we can not automatically determine the directions of all edges on the subgraph given the root. Our current implementation has two options. By default, we automatically find the minimum spanning tree of the subgraph of $\mathcal{B}$ on $\mathcal{A}$ into a tree by finding the minimum spanning tree of the subgraph and then subsquentially turn it into a directed tree. Alternatively, we ask the user to specify the direction of every edge through a `networkx.DiGraph` object.

The pseudotime for vertex $j$ is defined as:

$$o_j = \begin{cases} \max\left\{ \sum_{\ell \in \mathcal{L}} b_\ell : \mathcal{L} \subset \mathcal{E}(\overrightarrow{\mathcal{B}}) \text{ forms a simple path from } k_0 \text{ to } j \right\} \\ \infty \quad \text{if no such } \mathcal{L} \text{ exists} \end{cases} \tag{E.1}$$

where a simple path is a path that does not have any repeating vertices. Let $\boldsymbol{o} = (o_1, \cdots, o_k)$, then the pseudotime of the cell $i$ is defined as

$$T_i = \boldsymbol{o}^\top \tilde{\boldsymbol{w}}_i \tag{E.2}$$

which equals the pseudotime of the vertex that the cell is on or the weighted average of the pseudotime of the two vertices of the edge that the cell belongs to.

By our definition, any vertex that precedes another vertex on a simple path that starts from $k_0$ has a smaller pseudotime than the other vertex. Thus our definition guarantees a meaningful ordering of the cells when the directed trajectory backbone $\overrightarrow{\mathcal{B}}$ does not contain any cycles. If $\overrightarrow{\mathcal{B}}$ contains cycles, the pseudotime of the cells remains well-defined under our definition but may not be biologically meaningful.

Now, we relate the observed single-cell sequencing data with the underlying trajectory backbone graph $\mathcal{B}$. Let $\boldsymbol{Y}_i = (y_{i1}, \cdots, y_{iG})$ be the observed counts of $G$ features in cell $i$. Though $\boldsymbol{Y}_i$ is the vector of observed counts with complicated dependence across features, we assume that the dependence can be explained by latent Gaussian variables $\boldsymbol{Z}_i \in \mathbb{R}^d$, which is in a space with a much lower dimension and associated with the trajectory backbone $\mathcal{B}$. We also take into account the effects of known confounding covariates $\boldsymbol{X}_i \in \mathbb{R}^s$ (such as cell-cycle or batch effects, which we take as deterministic variables) on $\boldsymbol{Y}_i$. Specifically, we assume the following latent variable model on the observed counts:

$$\begin{aligned} \boldsymbol{Z}_i | \tilde{\boldsymbol{w}}_i &\sim \mathcal{N}_d(\boldsymbol{U}\tilde{\boldsymbol{w}}_i, \boldsymbol{I}_d) \\ Y_{ig} | \boldsymbol{Z}_i, \boldsymbol{X}_i &\overset{\text{ind}}{\sim} \mathcal{NB}\left(l_i f_g(\boldsymbol{Z}_i, \boldsymbol{X}_i), \theta_g\right), \quad g = 1, 2, \cdots, G \end{aligned} \tag{E.3}$$

Here $\boldsymbol{U} \in \mathbb{R}^{d \times k}$ is the unknown matrix of the vertices positions which, together with the cells' relative positions on $\mathcal{B}$, determines the means of $\boldsymbol{Z}_i$. Then, we assume that the observed counts $\boldsymbol{Y}_i$ nonlinearly depend on the latent variables $\boldsymbol{Z}_i$. The scalar $l_i$ is the known library size of cell $i$, and each $f_g : \mathbb{R}^{d+s} \longrightarrow \mathbb{R}, g = 1, 2, \cdots, G$ is an unknown nonlinear function involving the confounding covariate vector $\boldsymbol{X}_i$. The unknown parameters $\theta_g$ are feature-specific dispersion parameters of the Negative Binomial (NB) distribution. Notice that though the edges are assumed to be linear lines in the latent space, they are likely curves in the observed data space via the nonlinear mappings $\{f_g(\cdot),\ g = 1, 2, \cdots, G\}$.

The assumption of using NB distributions to model single-cell sequencing data is based on a detailed review and discussion in [45]. Specifically, for scRNA-seq with the unique molecular identifier (UMI) counts, NB distributions can describe the stochasticity in scRNA-seq, accounting for both biological and technical noise. However, they may not be adequate for sequencing data where the non-zero counts are large, such as scRNA-seq data without UMI. In that scenario, we assume that the observed counts follow zero-inflated

Negative Binomial (ZINB) distributions, as in [37]:

$$Y_{ig}|\boldsymbol{Z}_i, \boldsymbol{X}_i \overset{\text{ind}}{\sim} \phi_{ig}\delta_0 + (1-\phi_{ig})\mathcal{NB}\left(l_i f_g(\boldsymbol{Z}_i, \boldsymbol{X}_i), \theta_g\right) \tag{E.4}$$

where $\phi_{ig} = h_g(\boldsymbol{Z}_i, \boldsymbol{X}_i)$ is the zero inflation probability with $h_g(\cdot)$ an unknown non-linear function for $g = 1, \cdots, G$.

The above model defines the trajectory backbone $\mathcal{B}$, cell positions $\tilde{\boldsymbol{w}}_i$, and pseudotime $T_i$ that are identifiable from single-cell sequencing data. However, estimating these quantities is still challenging as the vertices and edges are defined on a latent space. Thus, we further impose a hierarchical model on $\tilde{\boldsymbol{w}}_i$ to simplify model estimation. First, we introduce a latent categorical variable $c_i$ as the index of all edges and vertices. Specifically, let $c_i$ take values in $\{1, 2, \ldots, K\}$ where $K = k(k+1)/2$ is the number of all possible edges and vertices in $\mathcal{G}$. For the complete graph $\mathcal{G}$, define the categorical assignment symmetric matrix,

$$\boldsymbol{C} = \begin{array}{c} \\ 1 \\ 2 \\ \vdots \\ k \end{array} \begin{array}{c} \begin{array}{cccc} 1 & 2 & \cdots & k \end{array} \\ \begin{pmatrix} 1 & 2 & \cdots & k \\ 2 & k+1 & \cdots & 2k-1 \\ \vdots & \vdots & \ddots & \vdots \\ k & 2k-1 & \cdots & K \end{pmatrix} \end{array}. \tag{E.5}$$

When $c_i$ equals to $C_{j_1 j_2}$ for $j_1 \neq j_2$, the cell $i$ is at the edge between vertex $j_1$ and vertex $j_2$. When $c_i$ equals to $C_{jj}$, the cell $i$ is at the vertex $j$. We let $C_{j_2 j_1} = C_{j_1 j_2} = j_1 + (j_2 - 1)k$ for $j_1 \leq j_2$ and set $c_i = C_{j_1 j_2}$ if cell $i$ is at the edge between vertex $j_1$ and vertex $j_2$, or if cell $i$ is at the vertex $j_1$ when $j_2 = j_1$. Then we assume the following mixture prior on $\tilde{\boldsymbol{w}}_i$:

$$\begin{aligned} w_i &\overset{\text{i.i.d.}}{\sim} \text{Uniform}(0, 1) \\ c_i &\overset{\text{i.i.d.}}{\sim} \text{Multinomial}(1, \boldsymbol{\pi}), \quad \boldsymbol{\pi} \in [0, 1]^K \\ w_i &\perp\!\!\!\perp c_i \\ \tilde{\boldsymbol{w}}_i &= w_i \boldsymbol{a}_{c_i} + (1 - w_i)\boldsymbol{b}_{c_i} \end{aligned} \tag{E.6}$$

where $\boldsymbol{a}_c = \boldsymbol{e}_{j_1}$ and $\boldsymbol{b}_c = \boldsymbol{e}_{j_2}$ if $c = C_{j_1 j_2}$ for $j_1 \leq j_2$.

To summarize, combining models (E.3) and (E.6), we obtain a hierarchical mixture model with an underlying trajectory structure. The unknown parameters are the mean positions of the $k$ cell types in the latent space $\boldsymbol{U}$, the prior probabilities of the $K$ categories $\boldsymbol{\pi}$, the mapping functions $f_g(\cdot), g = 1, 2, \cdots, G$ and the dispersion parameters $\theta_g$ (and also $h_g(\cdot)$ for the ZINB model).

**VAE for approximating the posterior distributions** To introduce a wide class of non-linear mapping functions, we model $f_g(\cdot)$ by a neural network and further combine our model with the variational autoencoder (VAE) [46] for approximating the posterior distributions. Following the variational Bayes approach in VAE and the conditional VAE [47], we use Gaussian distributions to approximate the intractable posterior distribution of $\boldsymbol{Z}_i$ given the observed data $\boldsymbol{Y}_i$ and confounding variables $\boldsymbol{X}_i$:

$$\boldsymbol{Z}_i|\boldsymbol{Y}_i, \boldsymbol{X}_i \dot{\sim} \mathcal{N}\left(\boldsymbol{\mu}_{\boldsymbol{Y}_i,\boldsymbol{X}_i}, \boldsymbol{\Sigma}_{\boldsymbol{Y}_i,\boldsymbol{X}_i}\right). \tag{E.7}$$

Here the "posterior" mean $\boldsymbol{\mu}_{\boldsymbol{Y}_i,\boldsymbol{X}_i}$ and covariance $\boldsymbol{\Sigma}_{\boldsymbol{Y}_i,\boldsymbol{X}_i} = \mathrm{diag}(\sigma^2_{\boldsymbol{Y}_i,\boldsymbol{X}_i,1}, \cdots, \sigma^2_{\boldsymbol{Y}_i,\boldsymbol{X}_i,d})$ are functions of $(\boldsymbol{Y}_i, \boldsymbol{X}_i)$. To guarantee flexibility, they are also modeled by a neural network. The neural network for $\boldsymbol{\mu}_{\boldsymbol{Y}_i,\boldsymbol{X}_i}$ and $\boldsymbol{\Sigma}_{\boldsymbol{Y}_i,\boldsymbol{X}_i}$ is the encoder, and the network for $f_g(\boldsymbol{Z}_i, \boldsymbol{X}_i)$ is the decoder. For non-UMI data, as in [37], the functions $h_g(\cdot)$ for zero-inflation parameters are also modeled in the decoder and share the same hidden layers as $f_g(\cdot)$.

With the above setup, we can lower bound the log-likelihood of each observation $\boldsymbol{Y}_i$ as

$$\begin{aligned}\log p(\boldsymbol{Y}_i|\boldsymbol{X}_i) &= \log\left[\mathbb{E}_{p(\boldsymbol{Z}_i|\boldsymbol{X}_i)} p(\boldsymbol{Y}_i|\boldsymbol{Z}_i, \boldsymbol{X}_i)\right] \\ &\geq \mathbb{E}_{q(\boldsymbol{Z}_i|\boldsymbol{Y}_i,\boldsymbol{X}_i)} \log p(\boldsymbol{Y}_i|\boldsymbol{Z}_i, \boldsymbol{X}_i) - D_{\mathrm{KL}}(q(\boldsymbol{Z}_i|\boldsymbol{Y}_i, \boldsymbol{X}_i)\|p(\boldsymbol{Z}_i)),\end{aligned} \tag{E.8}$$

where $p(\boldsymbol{Y}_i|\boldsymbol{Z}_i, \boldsymbol{X}_i)$ is the true conditional distribution in model (E.3) or (E.4), and $q(\boldsymbol{Z}_i|\boldsymbol{Y}_i, \boldsymbol{X}_i)$ is the Gaussian approximation of the posterior distribution in (E.7). This lower bound is often referred to as the evidence lower bound (ELBO) [46], where the first term $\mathbb{E}_{q(\boldsymbol{Z}_i|\boldsymbol{Y}_i,\boldsymbol{X}_i)} \log p(\boldsymbol{Y}_i|\boldsymbol{Z}_i, \boldsymbol{X}_i)$ denotes the reconstruction likelihood and the second term $-D_{\mathrm{KL}}(q(\boldsymbol{Z}_i|\boldsymbol{Y}_i, \boldsymbol{X}_i)\|p(\boldsymbol{Z}_i))$ behaves as a regularizer. The difference between (E.8) and regular conditional VAE is that we have a mixture model for the trajectory structure encoded in the term $p(\boldsymbol{Z}_i)$, which encourages the posteriors of $\boldsymbol{Z}_i$ to lie along linear edges and vertices. The resulting loss function for one cell is defined as the negative modified ELBOs in (E.8),

$$\mathcal{L}(\boldsymbol{Y}_i; \boldsymbol{X}_i, \Theta) = -\mathbb{E}_{q(\boldsymbol{Z}_i|\boldsymbol{Y}_i,\boldsymbol{X}_i)} \log p(\boldsymbol{Y}_i|\boldsymbol{Z}_i, \boldsymbol{X}_i) + \beta D_{\mathrm{KL}}(q(\boldsymbol{Z}_i|\boldsymbol{Y}_i, \boldsymbol{X}_i)\|p(\boldsymbol{Z}_i)), \tag{E.9}$$

where $\Theta$ represents all unknown parameters, including $\boldsymbol{U}$, $\boldsymbol{\pi}$, the unknown weights in the encoder and the decoder, and the dispersion parameters $\theta_g$. The tuning parameter $\beta$, introduced in [48] as the $\beta$-VAE, is to balance the reconstruction error and regularization. To better adjust for the confounding $\boldsymbol{X}_i$ and increase robustness, we add three extra (optional) penalties to obtain our final loss function aggregated over all $N$ cells as:

$$\begin{aligned}\mathcal{L}(\boldsymbol{Y}_1, \cdots, \boldsymbol{Y}_N; \boldsymbol{X}_1, \cdots \boldsymbol{X}_N, \Theta) = &- (1-\alpha)\sum_{i=1}^{N} \mathbb{E}_{q(\boldsymbol{Z}_i|\boldsymbol{Y}_i,\boldsymbol{X}_i)} \log p(\boldsymbol{Y}_i|\boldsymbol{Z}_i, \boldsymbol{X}_i) \\ &- \alpha \sum_{i=1}^{N} \log p(\boldsymbol{Y}_i|\boldsymbol{Z}_i = \boldsymbol{0}_d, \boldsymbol{X}_i) + \beta \sum_{i=1}^{N} D_{\mathrm{KL}}(q(\boldsymbol{Z}_i|\boldsymbol{Y}_i, \boldsymbol{X}_i)\|p(\boldsymbol{Z}_i)) \\ &+ \gamma\, \Omega_{\mathrm{Jacobian}}(\boldsymbol{Y}_1, \cdots, \boldsymbol{Y}_N; \boldsymbol{X}_1, \cdots \boldsymbol{X}_N) + \kappa\, \Omega_{\mathrm{MMD}}(\boldsymbol{Y}_1, \cdots, \boldsymbol{Y}_N; \boldsymbol{X}_1, \cdots \boldsymbol{X}_N).\end{aligned} \tag{E.10}$$

where $\alpha$ is a tuning parameter for "soft" batch adjustment [11], $\Omega_{\mathrm{MMD}}$ is the MMD loss to remove the difference of cells from different sources, and $\Omega_{\mathrm{Jacobian}}$ is the Jacobian regularizer to stabilize the optimization processes (see Appendix S.2.1 for more details). Setting $\alpha > 0$ ($\alpha = 0.1$ by default) encourages our decoder to reconstruct $\boldsymbol{Y}_i$ using only information from $\boldsymbol{X}_i$, which can help to decorrelate $\boldsymbol{X}_i$ from $\boldsymbol{Z}_i$ so that one can remove the confounding effects of $\boldsymbol{X}_i$ in $\boldsymbol{Z}_i$ more thoroughly. We set $\gamma = 1, \kappa = 0$ as default.

All the terms in (E.10) can be approximated by the Monte Carlo method efficiently, as we show in Appendix S.2.1. Specifically, though the marginal density function $p(\boldsymbol{Z}_i)$ is involved in a complex hierarchical mixture model (E.6), it still has a closed-form representation. The optimization minimizing the loss function (E.10) can be efficiently done via stochastic backpropagation [49] [50] on mini-batches of data, with the commonly used amortized variation inference [51] for VAE.

##### S.1.2 Trajectory Inference from Posterior Approximations

After the training step, the model returns the estimated parameters, including $\widehat{\boldsymbol{U}}$, $\widehat{\boldsymbol{\pi}}$, the encoder $\widehat{q}(\boldsymbol{Z}_i|\boldsymbol{Y}_i, \boldsymbol{X}_i)$, and the decoder $\widehat{f}(\boldsymbol{Z}_i|\boldsymbol{Y}_i, \boldsymbol{X}_i)$. Replacing the true posterior density $p(\boldsymbol{Z}_i|\boldsymbol{Y}_i, \boldsymbol{X}_i)$ with the approximate posterior density $\widehat{q}(\boldsymbol{Z}_i|\boldsymbol{Y}_i, \boldsymbol{X}_i)$, we can also obtain an approximation of the posterior distribution of $(w_i, c_i)$ for each cell. Specifically, let $\boldsymbol{z}_i^{(1)}, \cdots, \boldsymbol{z}_i^{(L)}$ be $L$ ($L = 300$ by default) random samples from $\widehat{q}(\boldsymbol{Z}_i|\boldsymbol{Y}_i, \boldsymbol{X}_i)$, and use the fact that $(w_i, c_i) \perp\!\!\!\perp (\boldsymbol{Y}_i, \boldsymbol{X}_i)|\boldsymbol{Z}_i$, we can approximate the posterior density of $(w_i, c_i)$ as

$$\widehat{p}(w, c|\boldsymbol{Y}_i, \boldsymbol{X}_i) = \frac{1}{L}\sum_{l=1}^{L} \widehat{p}(w, c|\boldsymbol{Z}_i = z_i^{(l)})$$

where $\widehat{p}(w, c|\boldsymbol{Z}_i)$ is obtained by plugging in $\widehat{\boldsymbol{U}}$ and $\widehat{\boldsymbol{\pi}}$ into the true posterior density of $(w_i, c_i)$ given $\boldsymbol{Z}_i$. Similarly, we can get the approximate posterior distributions of $c_i$, the edge or vertex the cell belongs to, as

$$\widehat{p}(c_i = c|\boldsymbol{Y}_i, \boldsymbol{X}_i) = \frac{1}{L}\sum_{l=1}^{L} \widehat{p}(c_i = c|\boldsymbol{Z}_i = z_i^{(l)})$$

For the cell position $\tilde{\boldsymbol{w}}_i$ on the graph $\mathcal{G}$, as it is a function of $w_i$ and $c_i$ as defined in (E.6), we can also efficiently obtain the mean $\boldsymbol{\mu}_{\tilde{\boldsymbol{w}}_i}$ and the diagonal elements of the covariance matrix $\Sigma_{\tilde{\boldsymbol{w}}_i}$ of its posterior distribution $\widehat{p}(\tilde{\boldsymbol{w}}_i|\boldsymbol{Y}_i, \boldsymbol{X}_i)$. For details in calculating the approximate posterior distributions, see Appendix S.2.2. Now, we discuss how to infer the trajectory backbone and cell positions along the trajectory with these posterior approximations.

**Infer the trajectory backbone $\mathcal{B}$** The total number of categories $K = O(k^2)$ can be large, even with a moderate number of $k$. While the trajectory backbone $\mathcal{B}$ typically only sparsely involves a few edges, the estimated $\widehat{\boldsymbol{\pi}}$ may be dense on the unpruned entries. Inspired by [52] on Bayesian Gaussian mixture models, to encourage sparsity, we infer the nonzero edges a posteriori from the data. Specifically, we define a score for each edge, quantifying the strength of the evidence that the edge exists:

$$s_{j_1 j_2} = \frac{|\{i : c_i = C_{j_1 j_2}\}|}{|\{i : \boldsymbol{e}_{j_1}^\top \tilde{\boldsymbol{w}}_i > 0.5 \text{ or } \boldsymbol{e}_{j_2}^\top \tilde{\boldsymbol{w}}_i > 0.5\}|}.$$

where the denominator is added to make sure that we can capture the continuous transitions between cell states even when these states only involve a small proportion of cells in the cell population. Even though $\widehat{\boldsymbol{\pi}}$ is dense, $s_{j_1 j_2}$ can be significantly nonzero for much fewer edges. From another point of view, an edge score is some "test statistics" to evaluate whether the edge exists. To make our algorithm scalable, in practice, instead of obtaining the posterior distributions of $s_{j_1 j_2}$ to determine whether it is significantly nonzero or not, we simply use a deterministic version of the edge score as

$$\tilde{s}_{j_1 j_2} = \frac{|\{i : \tilde{c}_i = C_{j_1 j_2}\}|}{|\{i : \boldsymbol{e}_{j_1}^\top \boldsymbol{\mu}_{\tilde{\boldsymbol{w}}_i} > 0.5 \text{ or } \boldsymbol{e}_{j_2}^\top \boldsymbol{\mu}_{\tilde{\boldsymbol{w}}_i} > 0.5\}|}$$

where $\tilde{c}_i = \arg\max_{c \in \{1,2,\cdots,K\}} \widehat{p}(c_i = c | \boldsymbol{Y}_i, \boldsymbol{X}_i)$ and $\boldsymbol{\mu}_{\tilde{\boldsymbol{w}}_i}$ is the approximate posterior mean of $\tilde{\boldsymbol{w}}_i$. Other edge score choices are discussed in Appendix S.2.3.

A larger $\tilde{s}_{j_1 j_2}$ indicates higher confidence assuring that the edge exits in the trajectory backbone $\mathcal{B}$. In practice, we include an edge $(j_1, j_2)$ into the estimated backbone $\widehat{\mathcal{B}}$ if $\tilde{s}_{j_1 j_2} \geq s_0$, where the cutoff $s_0$ is 0.01 by default. When we are certain that there are no loops in the trajectory, inspired by [3], we further prune $\widehat{\mathcal{B}}$ as the MST of the unpruned graph with $\tilde{s}_{j_1 j_2}$ as edge weights. This typically results in a cleaner shape of our estimated trajectory.

**Project cells onto the inferred trajectory backbone** To obtain the position of each cell on the inferred trajectory, we further project $\tilde{\boldsymbol{w}}_i$ onto $\widehat{\mathcal{B}}$. Given the approximate posterior distributions of $\tilde{\boldsymbol{w}}_i$, we would find a point estimate for the best position of each cell $i$ on $\widehat{\mathcal{B}}$. Specifically, for each cell, we aim to solve the following optimization problem:

$$\begin{aligned} \widehat{\tilde{\boldsymbol{w}}}_i = \arg\min_{\boldsymbol{w}} \quad & \mathbb{E}_{\widehat{p}(\tilde{\boldsymbol{w}}_i | \boldsymbol{Y}_i, \boldsymbol{X}_i)} \|\tilde{\boldsymbol{w}}_i - \boldsymbol{w}\|_2^2 \\ \text{s.t.} \quad & \text{support}(\boldsymbol{w}) \subseteq \widehat{\mathcal{B}},\ \|\boldsymbol{w}\|_1 = 1,\ \boldsymbol{w} \succeq \boldsymbol{0}. \end{aligned} \tag{E.11}$$

Since the support of $\tilde{\boldsymbol{w}}_i$ is restricted to the inferred trajectory backbone $\widehat{\mathcal{B}}$, only one or two entries of $\widehat{\tilde{\boldsymbol{w}}}_i$ can be nonzero, depending on whether the cell is at the vertex or edge of $\widehat{\mathcal{B}}$. Though the $L_2$ loss in the objective function of optimization problem (E.11) is not the only choice, it can result in a closed-form solution that allows fast computation.

**Proposition 1.** *The optimization problem (E.11) is equivalent to finding*

$$\begin{aligned} \widehat{\tilde{\boldsymbol{w}}}_i = \arg\min_{\boldsymbol{w}} \quad & \|\boldsymbol{\mu}_{\tilde{\boldsymbol{w}}_i} - \boldsymbol{w}\|_2^2 \\ \text{s.t.} \quad & \text{support}(\boldsymbol{w}) \subseteq \widehat{\mathcal{B}},\ \|\boldsymbol{w}\|_1 = 1,\ \boldsymbol{w} \succeq \boldsymbol{0}, \end{aligned} \tag{E.12}$$

*where $\boldsymbol{\mu}_{\tilde{\boldsymbol{w}}_i}$ is the mean of $\widehat{p}(\tilde{\boldsymbol{w}}_i | \boldsymbol{Y}_i, \boldsymbol{X}_i)$. Denote the $j$th component of $\boldsymbol{\mu}_{\tilde{\boldsymbol{w}}_i}$ as $\mu_j$ and let $\mathcal{EN}(\widehat{\mathcal{B}}) = \mathcal{E}(\widehat{\mathcal{B}}) \cup \left\{(j,j) : j \in \mathcal{N}(\widehat{\mathcal{B}})\right\}$, then the best projection is given by*

$$(j_1^*, j_2^*) \in \underset{(j_1,j_2) \in \mathcal{EN}(\widehat{\mathcal{B}})}{\arg\max} \quad (\mu_{j_1} - \mu_{j_2})^2 + 2(\mu_{j_1} + \mu_{j_2}) - \mathbb{1}_{\{j_1 = j_2\}},$$

*and the corresponding solution* $\text{proj}_{(j_1^*,j_2^*)}(\boldsymbol{\mu}_{\tilde{\boldsymbol{w}}_i})$ *has entries*

$$[\text{proj}_{(j_1^*,j_2^*)}(\boldsymbol{\mu}_{\tilde{\boldsymbol{w}}_i})]_j = \begin{cases} \mu_{j_1^*} + \frac{1}{2}\left(1 - \mu_{j_1^*} - \mu_{j_2^*} + \mathbb{1}_{\{j_1^*=j_2^*\}}\right) & j \in \{j_1^*, j_2^*\} \\ 0 & \text{otherwise,} \end{cases}$$

*for* $j = 1, \ldots, k$. *The best projection reduces to be a vertex if* $j_1^* = j_2^*$.

The proof of Proposition 1 is included in Appendix S.2.4. Intuitively, finding the optimal projection of $\tilde{\boldsymbol{w}}_i$ minimizing the $L_2$ loss is equivalent to simply projecting the posterior mean $\boldsymbol{\mu}_{\tilde{\boldsymbol{w}}_i}$. In addition, as shown in our proof of Proposition 1, most cells will either project onto an edge or an isolated vertex. Notice that the solution for the optimization problem (E.12) may not be unique. However, it is generally unique due to floating-point computation in practice.

Next, we also want to quantify the uncertainty of the projected position $\widehat{\boldsymbol{w}}_i$ as an estimate of $\tilde{\boldsymbol{w}}_i$. A general metric to evaluate the uncertainty is given by $\mathbb{E}_{\widehat{p}(\tilde{\boldsymbol{w}}_i|\boldsymbol{Y}_i,\boldsymbol{X}_i)}[d(\widehat{\boldsymbol{w}}_i, \tilde{\boldsymbol{w}}_i)]$, where $d(\cdot,\cdot)$ is a metric or distance function. When $d(\boldsymbol{w}, \boldsymbol{w}') = \|\boldsymbol{w} - \boldsymbol{w}'\|_2^2$ is also the $L_2$ loss, we obtain the projection mean square error (MSE) as

$$\mathbb{E}_{\widehat{p}(\tilde{\boldsymbol{w}}_i|\boldsymbol{Y}_i,\boldsymbol{X}_i)}[d(\widehat{\boldsymbol{w}}_i, \tilde{\boldsymbol{w}}_i)] = \|\widehat{\boldsymbol{w}}_i - \boldsymbol{\mu}_{\tilde{\boldsymbol{w}}_i}\|_2^2 + \text{tr}\left(\Sigma_{\tilde{\boldsymbol{w}}_i}\right), \tag{E.13}$$

which is easily computable. Notice that our projection MSE ignores the uncertainty in $\widehat{\mathcal{B}}$ and the approximation error in $\widehat{p}(\tilde{\boldsymbol{w}}_i|\boldsymbol{Y}_i, \boldsymbol{X}_i)$, so it is an underestimate of the true uncertainty. However, we think that the relative magnitude of our projection MSE would still be a useful quantity to compare the projection accuracy across cells. Another pattern is that the projection MSE is typically smaller when the cells are near vertices. We provide our understanding and a detailed discussion of this pattern in Appendix S.1.5.

Finally, we obtain the point estimate of each cell's pseudotime $T_i$ defined in (7). Given the inferred trajectory $\widehat{\mathcal{B}}$, the user can assign a root vertex based on prior biological knowledge. We also provide an automatic root selection step following Tempora [39] when cells are collected from a series of time points. The idea is to choose the vertex with the earliest collection time as the root. Specifically, let $r_i$ be the collection time of cell $i$, we calculate the "collection time" of vertex $j$ as $\omega_j = \sum_i \widehat{\boldsymbol{w}}_{ij} r_i / \sum_i \widehat{\boldsymbol{w}}_{ij}$, a weighted average of the collection time of the cells near the vertex. A vertex $k_0 \in \mathcal{N}(\mathcal{B})$ is chosen as the root if it has the smallest collection time $\omega_j$ and is not an isolated vertex.

Once the root vertex of the trajectory backbone $\widehat{\mathcal{B}}$ is obtained, it is straightforward to obtain $\widehat{\boldsymbol{o}}$, the estimated pseudotime of the vertices, by plugging in $\widehat{\mathcal{B}}$ into the definition of $\boldsymbol{o}$ in (E.1). Then, a point estimate of the pseudotime $T_i$ of cell $i$ can be given as $\widehat{T}_i = \widehat{\boldsymbol{o}}^\top \widehat{\boldsymbol{w}}_i$.

##### S.1.3 Differential gene expression along the trajectory

We provide a polynomial regression approach to find differentially expressed genes along our inferred trajectory backbone. We focus on finding genes that are associated with the pseudotime ordering after adjusting for confounding covariates and provide a scalable way to obtain the $p$-values of the genes, taking into consideration that the pseudotimes of the cells are estimated.

In particular, to find genes that are differentially expressed along the pseudotime ordering for a subset of cells $\mathcal{S}$, we work with the following polynomial regression for each gene $g$:

$$Y_{ig} = \beta_{0g} + \beta_{1g}\text{rank}(T_i) + \cdots + \beta_{Kg}\text{rank}(T_i)^K + \boldsymbol{X}_i^\top \boldsymbol{\beta}_{Xg} + e_{ig}, \quad \mathbb{E}(e_{ig}) = 0, \ \forall\, i \in \mathcal{S} \tag{E.14}$$

where $Y_{ig}$ is the log-transformed and normalized count for cell $i$ and gene $g$, $\text{rank}(T_i)$ is the rank of true pseudotime $T_i$ of cell $i$, and the linear term $\boldsymbol{X}_i^\top \boldsymbol{\beta}_{Xg}$ is to adjust for the confounding effects of known covariates. We allow $e_{ig}$'s to have unequal variances as $Y_{ig}$'s are log-transformed and normalized counts. $K$ is the degree of the polynomials and by default, is chosen as 2 to allow both linear and quadratic change of the gene expression along pseudotimes. For each gene, we aim to test for the global null $H_{0g} : \beta_{1g} = \cdots = \beta_{Kg} = 0$. We use $\text{rank}(T_i)$ in the regression model so that the results will not be affected by the scaling of pseudotimes.

The challenge here is that the true pseudotime $T_i$ is not observed. Instead, we only have estimated pseudotime $\widehat{T}_i$ from the data, which has unknown uncertainty and is correlated with $Y_{ig}$. As a consequence, the $p$-value for $H_{0g}$ ignoring the fact that the pseudotimes of the cells are estimated is likely invalid. To adjust for this, we take a simple p-value calibration approach following [40]. To get an initial p-value, we plug in $\widehat{T}_i$ after normalizing and centering each $\text{rank}(T_i)^k$, and we estimate the coefficients $\beta_{kg}$ via the ordinary least squares. Treating $\widehat{T}_i$ as the true $T_i$, we obtain the variances of $\widehat{\beta}_{kg}$ through the sandwich estimator, allowing for heterogeneity in $e_{ig}$. The $t$-statistics is then defined as $t_{kg} = \widehat{\beta}_{kg}/\widehat{\text{sd}}(\widehat{\beta}_{kg})$. To calibrate the p-values, instead of assuming $t_{kg} \sim \mathcal{N}(0, 1)$ under the null, we assume that $t_{kg} \sim \mathcal{N}(0, \sigma_k^2)$ where $\sigma_k^2$ is estimated from the median absolute deviation (MAD) of $\{t_{kg},\ g = 1, 2, \cdots, G\}$. Then we use the Bonferroni combination approach across $k$ to obtain a combined $p$-value for the null $H_{0g}$. To select differentially expressed genes, these $p$-values are further adjusted with the Benjamini-Hochberg procedure for multiple testing corrections.

There are other approaches to detecting differentially expressed genes along estimated pseudotime. For instance, tradeSeq [16] employs a generalized linear regression model on raw scRNA-seq counts, assuming a Negative Binomial distribution of the data. They regress $Y_{ig}$ on the smoothing splines of $\widehat{T}_i$. PseudotimeDE [17] uses a sophisticated approach to calibrate the p-values in differential testing with estimated pseudotime. To adjust for the bias treating $\widehat{T}_i$ as the true pseudotime, PseudotimeDE employs cell subsampling and permutation of estimated pseudotime to generate a null distribution for the regression test statistics of each gene. Compared to tradeSeq and pseudotimeDE, our approach sacrifices some rigor for increased computational efficiency. See Appendix S.3.3 for an empirical comparison of our differential testing approach with PseudotimeDE and tradeSeq.

##### S.1.4 Model estimation with practical considerations

Instead of using all the genes, by default, we select highly variable genes and preprocess the gene expressions by scanpy [42]. Then the normalized, log-transformed, and scaled gene expressions are provided as the inputs of our VAE.

As the optimization of our loss (E.10) generally results in a locally optimal solution, we need a good initialization of our parameters, especially for $\boldsymbol{U}$ and $\boldsymbol{\pi}$ defined in (E.3) and (E.6). Also, our framework requires a pre-determined number of states $k$. Inspired by other existing TI methods, we design a three-step algorithm for model initialization and estimation.

The first step is pretraining, where we train the model with $\beta = 0$, to only minimize the reconstruction loss, which does not involve the unknown parameters $\boldsymbol{\pi}$ and $\boldsymbol{U}$. The Jacobian and MMD regularizers can also be used in this stage. The purpose of this step is to obtain better weights for the encoder and the decoder and to get an initial low-dimensional representation, which can be used to initialize $\boldsymbol{U}$ and determine $k$.

The second step is to initialize the latent space after pretraining. In our experiments, we use annotated cell types to initialize the latent space and set $k$ to be the number of cell types. However, one can also perform cell clustering with the Louvain algorithm [53] on the estimated posterior means $\widehat{\boldsymbol{\mu}}_{\boldsymbol{Y}_i,\boldsymbol{X}_i}$ of $\boldsymbol{Z}_i$ to determine $k$. We initialize $\boldsymbol{U}$ with the cluster centers. As $\boldsymbol{\pi}$ involves both the $k$ vertices and $k(k-1)/2$ edges, we have no information yet and just uniformly initialize $\boldsymbol{\pi}$ in this step. We introduce a pruning mechanism for better estimating $\boldsymbol{\pi}$. Specifically, we set those $\pi_i$'s corresponding to edges with the top $50\%$ lengths (the distance between the initial centers) as 0 and freeze them when training.

The last step is to train our whole network, optimizing the loss function (E.10) with $\beta = 1$. During both the pretraining and training steps, the optimization is early stopped when the evaluation loss decreases sufficiently slowly.

Finally, to make VITAE scalable to handle large datasets and have a comparable computational cost as other TI methods, we accelerate VITAE for large datasets by reducing the dimension of the input. We replace the $G$ features with its top $R$ ($R = 64$ by default) principle component (PC) scores $\boldsymbol{F}_i$. The output of the decoder is also replaced by the reconstruction of $\boldsymbol{F}_i$, and the likelihood of $p(\boldsymbol{Y}_i|\boldsymbol{Z}_i, \boldsymbol{X}_i)$ in the loss function (E.10) is replaced by Gaussian densities assumed on $\boldsymbol{F}_i$ (Appendix S.2.1). As we will show in the Result Section, the accelerated VITAE with Gaussian densities provides results comparable to our original likelihood-based VITAE in general, while it can be much faster than the original scheme. By default, both the encoder and decoder have one hidden layer, with 32 and 16 units for the accelerated and likelihood-based VITAE, respectively. The bottleneck layer only has a dimension of 8. The hidden layers are all fully connected with the leaky rectified linear activation function [54] and Batch Normalization [55].

##### S.1.5 Discussion on Uncertainty Quantification

We observed in praactice that our calculated projection MSE is always smaller for the cells near vertices. Why do we observe such a pattern? Is it due to the estimation bias in our approximation of the posterior distribution, or is it an intrinsic property of the L2 loss under our mixture model? In order to answer these questions, we consider the case where we observe the latent space $\boldsymbol{Z}_i$, so that we can compute the true posterior distributions of $\boldsymbol{Z}_i|\tilde{\boldsymbol{w}}_i$. Also, we focus on $\mathrm{tr}(\Sigma_{\tilde{\boldsymbol{w}}_i})$, as from our observations it is typically the leading term in (12). Specifically, we define our hierarchical model with observed $\boldsymbol{Z}_i$ following (3) and (5) by setting $d = 2$, $k = 3$ with $\boldsymbol{U} = \begin{pmatrix} 0 & 0.5 & 2 \\ 2 & 0.5 & 0 \end{pmatrix}$. The defined backbone is shown in Figure S0.1a and we choose vertex 1 as the root of the trajectory.

First, we take $\boldsymbol{\pi} = \begin{pmatrix} \frac{1}{5} & \frac{1}{5} & 0 & \frac{1}{5} & \frac{1}{5} & \frac{1}{5} \end{pmatrix}$, so that the edges and vertices have equal probabilities. As $\boldsymbol{Z}_i$ is observed, we can compute the true posterior mean and variances from Appendix S.2.2 for any given $\boldsymbol{Z}_i$. Here we show the posterior variances given $\boldsymbol{Z}_i = \boldsymbol{U}\tilde{\boldsymbol{w}}_i$ which are exactly on the trajectory, and discuss how the variances change with $T_i = \boldsymbol{o}^\top \tilde{\boldsymbol{w}}_i$. The cell $i$ is at the vertices when $T_i = 0, 1$ or 2. Figure S0.1b shows how the posterior variances of each $\tilde{w}_{ij}$ change with $T_i$. The leading term of the projection MSE tr$(\Sigma_{\tilde{\boldsymbol{w}}_i})$ summing up the three terms in Figure S0.1b has an M-shape as shown in Figure S0.1c, indicating that the cells indeed have smaller projection uncertainties when they are near the vertices.

One explanation of the above phenomenon is that as there are nonzero probabilities exactly at the vertices, the distribution is denser near the vertices. As a consequence, there is less uncertainty on $\tilde{\boldsymbol{w}}_i$ if it is closer to a vertex. However, what we find surprising is that when we set $\boldsymbol{\pi} = \begin{pmatrix} 0 & \frac{1}{2} & 0 & 0 & \frac{1}{2} & 0 \end{pmatrix}$, where the cells can only be on the edges, we still observe an M-shape for the change of tr$(\Sigma_{\tilde{\boldsymbol{w}}_i})$ along $T_i$. In other words, the projection MSE from true posteriors is smaller near the vertices no matter whether the vertices have nonzero probabilities or not, though the difference is smaller when the vertices have zero probabilities. As a consequence, we believe that this pattern is an intrinsic property of the $L_2$ loss under our mixture model.

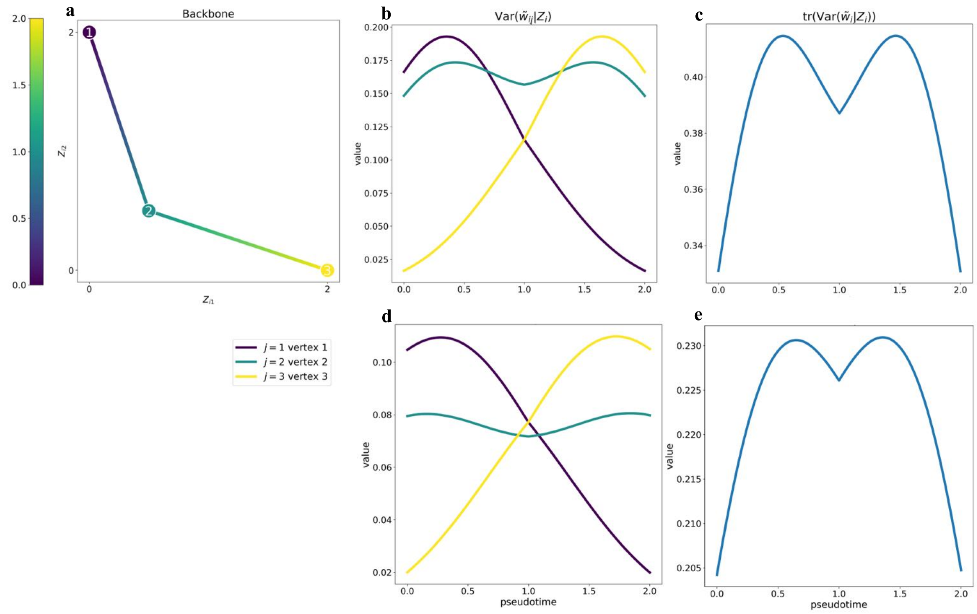

**Fig S0.1:** **(a)** The underlying backbone of the assumed model colored by pseudotime. **(b)** The values of Var$(\tilde{w}_{ij}|\boldsymbol{Z}_i)$ along the trajectory and **(c)** the values of tr(Var$(\tilde{\boldsymbol{w}}_i|\boldsymbol{Z}_i)$) along the trajectory are for the case when $\boldsymbol{\pi} = (0.2, 0.2, 0, 0.2, 0.2, 0.2)$. **(d-e)** are the same as **(b-c)** for the case when $\boldsymbol{\pi} = (0, 0.5, 0, 0, 0.5, 0)$.

#### S.2 Technical details

##### S.2.1 Loss Function

**Basic loss function.** Note that the loss for each cell in (E.10) takes the form

$$\begin{aligned}\mathcal{L}(\boldsymbol{Y}_i;\boldsymbol{X}_i,\Theta) &= -(1-\alpha)\mathbb{E}_{q(\boldsymbol{Z}_i|\boldsymbol{Y}_i,\boldsymbol{X}_i)}\log p(\boldsymbol{Y}_i|\boldsymbol{Z}_i,\boldsymbol{X}_i)\\ &\quad -\alpha\log p(\boldsymbol{Y}_i|\boldsymbol{Z}_i=\boldsymbol{0}_d,\boldsymbol{X}_i)+\beta D_{\mathrm{KL}}(q(\boldsymbol{Z}_i|\boldsymbol{Y}_i,\boldsymbol{X}_i)\|p(\boldsymbol{Z}_i))\\ &= -(1-\alpha)\mathbb{E}_{q(\boldsymbol{Z}_i|\boldsymbol{Y}_i,\boldsymbol{X}_i)}\log p(\boldsymbol{Y}_i|\boldsymbol{Z}_i,\boldsymbol{X}_i)-\alpha\log p(\boldsymbol{Y}_i|\boldsymbol{Z}_i=\boldsymbol{0}_d,\boldsymbol{X}_i)\\ &\quad +\beta\mathbb{E}_{q(\boldsymbol{Z}_i|\boldsymbol{Y}_i,\boldsymbol{X}_i)}\log q(\boldsymbol{Z}_i|\boldsymbol{Y}_i,\boldsymbol{X}_i)-\beta\mathbb{E}_{q(\boldsymbol{Z}_i|\boldsymbol{Y}_i,\boldsymbol{X}_i)}\log p(\boldsymbol{Z}_i),\end{aligned}$$

when $\Omega_{\text{MMD}}\equiv\Omega_{\text{Jacobian}}\equiv 0$. Next, we show how to compute the above terms efficiently.

1. $\mathbb{E}_{q(\boldsymbol{Z}_i|\boldsymbol{Y}_i,\boldsymbol{X}_i)}\log p(\boldsymbol{Y}_i|\boldsymbol{Z}_i,\boldsymbol{X}_i)$

$$\mathbb{E}_{q(\boldsymbol{Z}_i|\boldsymbol{Y}_i,\boldsymbol{X}_i)}\log p(\boldsymbol{Y}_i|\boldsymbol{Z}_i,\boldsymbol{X}_i)\approx\frac{1}{L}\sum_{l=1}^{L}\sum_{g=1}^{G}\log p(Y_{ig}|\boldsymbol{Z}_i=\boldsymbol{z}_i^{(l)},\boldsymbol{X}_i)$$

where $\boldsymbol{z}_i^{(1)},\ldots,\boldsymbol{z}_i^{(L)}$ are Monte Carlo samples from the approximate posterior distribution $q(\boldsymbol{Z}_i|\boldsymbol{Y}_i,\boldsymbol{X}_i)$. Here $p(Y_{ig}|\boldsymbol{Z}_i,\boldsymbol{X}_i)$ depends on the distribution assumptions of $Y_{ig}$.

(1) For scRNA-seq with UMI, $Y_{ig}|\boldsymbol{Z}_i,\boldsymbol{X}_i$ follows a Negative Binomial distribution $\mathcal{NB}(\lambda_{ig},\theta_g)$ with a probability mass function,

$$p(y|\boldsymbol{Z}_i,\boldsymbol{X}_i)=\binom{\theta_g+y-1}{y}\left(\frac{\theta_g}{\lambda_{ig}+\theta_g}\right)^{\theta_g}\left(\frac{\lambda_{ig}}{\lambda_{ig}+\theta_g}\right)^{y},$$

for $i=1,\ldots,N$. It is noted that only $\lambda_{ig}$ dependent on $\boldsymbol{Z}_i$ and $\boldsymbol{X}_i$.

(2) For non-UMI data, $Y_{ig}|\boldsymbol{Z}_i,\boldsymbol{X}_i$ follows a zero-inflated Negative Binomial distribution $\mathcal{ZINB}(\lambda_{ig},\theta_g,\phi_{ig})$ with the probability mass function,

$$p(y|\boldsymbol{Z}_i,\boldsymbol{X}_i)=\begin{cases}\phi_{ig}+(1-\phi_{ig})\,p_{NB}(0;\lambda_{ig},\theta_g) & \text{if } y=0\\ (1-\phi_{ig})\,p_{NB}(y;\lambda_{ig},\theta_g) & \text{if } y>0,\end{cases}$$

where $p_{NB}(y;\lambda_{ig},\theta_g)$ is the probability mass function of $\mathcal{NB}(\lambda_{ig},\theta_g)$.

(3) For the accelerated VITAE with a Gaussian model, $\boldsymbol{Y}_i$ is replaced by the PC scores $\boldsymbol{F}_i$. We assume that each $F_{ig}|\boldsymbol{Z}_i,\boldsymbol{X}_i$ follows a Gaussian distribution $\mathcal{N}(\nu_{ig},\tau_i^2)$ with probability density function,

$$p(y|\boldsymbol{Z}_i,\boldsymbol{X}_i)=\frac{1}{\sqrt{2\pi}\tau_i}\exp\left(-\frac{(y-\nu_{ig})^2}{2\tau_i^2}\right).$$

2. $\mathbb{E}_{q(\boldsymbol{Z}_i|\boldsymbol{Y}_i,\boldsymbol{X}_i)} \log p(\boldsymbol{Z}_i)$

$$\mathbb{E}_{q(\boldsymbol{Z}_i|\boldsymbol{Y}_i,\boldsymbol{X}_i)} \log p(\boldsymbol{Z}_i) = \mathbb{E}_{q(\boldsymbol{Z}_i|\boldsymbol{Y}_i,\boldsymbol{X}_i)} \left[ \log \left( \sum_{c=1}^{K} p(\boldsymbol{z}|c_i = c)p(c) \right) \right] \approx \frac{1}{L} \sum_{l=1}^{L} \log \left( \sum_{c=1}^{K} p(\boldsymbol{z}^{(l)}|c_i = c)p(c) \right).$$

When $c \in \mathcal{N}(\mathcal{G})$, the conditional density of $\boldsymbol{Z}_i|c_i = c$ is given by

$$p(\boldsymbol{z}|c_i = c) = \int_0^1 p(\boldsymbol{z}|c_i = c, w)\mathrm{d}w = \frac{1}{(2\pi)^{\frac{d}{2}}} \int_0^1 e^{-\frac{1}{2}(\boldsymbol{z}-\boldsymbol{U}\boldsymbol{b}_c)^\top(\boldsymbol{z}-\boldsymbol{U}\boldsymbol{b}_c)}\mathrm{d}w = \varphi_d(\boldsymbol{U}\boldsymbol{b}_c).$$

where $\varphi_d(\cdot)$ is the probability density function of the standard $d$-dimensional multivariate Gaussian distribution.

When $c \in \mathcal{E}(\mathcal{G})$, the conditional density of $\boldsymbol{Z}_i|c_i = c$ is given by

$$\begin{aligned} p(\boldsymbol{z}|c_i = c) &= \int_0^1 p(\boldsymbol{z}|c_i = c, w)\mathrm{d}w \\ &= \frac{1}{(2\pi)^{\frac{d}{2}}} \int_0^1 e^{-\frac{1}{2}(\boldsymbol{z}-\boldsymbol{U}\tilde{w})^\top(\boldsymbol{z}-\boldsymbol{U}\tilde{w})}\mathrm{d}w \\ &\overset{\substack{\boldsymbol{\alpha}_{\boldsymbol{z}c}=\boldsymbol{U}(\boldsymbol{b}_c-\boldsymbol{a}_c) \\ \boldsymbol{\beta}_{\boldsymbol{z}c}=\boldsymbol{z}-\boldsymbol{U}\boldsymbol{b}_c}}{=\!=\!=} \frac{1}{(2\pi)^{\frac{d}{2}}} \int_0^1 e^{-\frac{1}{2}(w\boldsymbol{\alpha}_{\boldsymbol{z}c}+\boldsymbol{\beta}_{\boldsymbol{z}c})^\top(w\boldsymbol{\alpha}_{\boldsymbol{z}c}+\boldsymbol{\beta}_{\boldsymbol{z}c})}\mathrm{d}w \\ &= \frac{1}{(2\pi)^{\frac{d}{2}}} \int_0^1 e^{-\frac{1}{2}(\boldsymbol{\alpha}_{\boldsymbol{z}c}^\top\boldsymbol{\alpha}_{\boldsymbol{z}c}w^2+2\boldsymbol{\alpha}_{\boldsymbol{z}c}^\top\boldsymbol{\beta}_{\boldsymbol{z}c}w+\boldsymbol{\beta}_{\boldsymbol{z}c}^\top\boldsymbol{\beta}_{\boldsymbol{z}c})}\mathrm{d}w \\ &= \frac{\sigma_c e^{\frac{t_c}{2}}}{(2\pi)^{\frac{d-1}{2}}} \left[ \Phi\left(\frac{1-\nu_{\boldsymbol{z}c}}{\sigma_c}\right) - \Phi\left(-\frac{\nu_{\boldsymbol{z}c}}{\sigma_c}\right) \right], \end{aligned}$$

with

$$\nu_{\boldsymbol{z}c} = -\frac{\boldsymbol{\alpha}_{\boldsymbol{z}c}^\top\boldsymbol{\beta}_{\boldsymbol{z}c}}{\boldsymbol{\alpha}_{\boldsymbol{z}c}^\top\boldsymbol{\alpha}_{\boldsymbol{z}c}}, \quad \sigma_{\boldsymbol{z}c}^2 = \frac{1}{\boldsymbol{\alpha}_{\boldsymbol{z}c}^\top\boldsymbol{\alpha}_{\boldsymbol{z}c}}, \quad t_{\boldsymbol{z}c} = -\boldsymbol{\beta}_{\boldsymbol{z}c}^\top\boldsymbol{\beta}_{\boldsymbol{z}c} + \frac{(\boldsymbol{\alpha}_{\boldsymbol{z}c}^\top\boldsymbol{\beta}_{\boldsymbol{z}c})^2}{\boldsymbol{\alpha}_{\boldsymbol{z}c}^\top\boldsymbol{\alpha}_{\boldsymbol{z}c}}$$

where $\Phi(\cdot)$ is the cumulative density function of the standard Gaussian distribution.

Therefore, the marginal density of $\boldsymbol{Z}_i$ is given by

$$p(\boldsymbol{z}) = \sum_{c=1}^{K} p(\boldsymbol{z}|c_i = c)\pi_c = \sum_{c\in\mathcal{N}(\mathcal{G})} \pi_c\varphi_d(\boldsymbol{U}\boldsymbol{b}_c) + \sum_{c\in\mathcal{E}(\mathcal{G})} \frac{\pi_c\sigma_{\boldsymbol{z}c}e^{\frac{t_{\boldsymbol{z}c}}{2}}}{(2\pi)^{\frac{d-1}{2}}} \left[ \Phi\left(\frac{1-\nu_{\boldsymbol{z}c}}{\sigma_{\boldsymbol{z}c}}\right) - \Phi\left(-\frac{\nu_{\boldsymbol{z}c}}{\sigma_{\boldsymbol{z}c}}\right) \right].$$

3. $\mathbb{E}_{q(\boldsymbol{Z}_i|\boldsymbol{Y}_i,\boldsymbol{X}_i)} \log q(\boldsymbol{Z}_i|\boldsymbol{Y}_i, \boldsymbol{X}_i)$

Since the approximate posterior density $q(\boldsymbol{Z}_i|\boldsymbol{Y}_i, \boldsymbol{X}_i)$ is $\mathcal{N}(\boldsymbol{\mu}_{\boldsymbol{Y}_i,\boldsymbol{X}_i}, \boldsymbol{\Sigma}_{\boldsymbol{Y}_i,\boldsymbol{X}_i})$, we have

$$\mathbb{E}_{q(\boldsymbol{Z}_i|\boldsymbol{Y}_i,\boldsymbol{X}_i)} \log q(\boldsymbol{Z}_i|\boldsymbol{Y}_i, \boldsymbol{X}_i)$$

$$
\begin{aligned}
=&\mathbb{E}_{q(\boldsymbol{Z}_i|\boldsymbol{Y}_i,\boldsymbol{X}_i)}\left[-\frac{d}{2}\log(2\pi)-\frac{1}{2}\sum_{i=1}^{d}\log\sigma^2_{\boldsymbol{Y}_i,\boldsymbol{X}_i,i}-\frac{1}{2}\sum_{j=1}^{d}\frac{(Z_{ij}-\mu_{\boldsymbol{Y}_i,\boldsymbol{X}_i,j})^2}{\sigma^2_{\boldsymbol{Y}_i,\boldsymbol{X}_i,j}}\right]\\
=&-\frac{d}{2}\log(2\pi)-\frac{1}{2}\sum_{i=1}^{d}\log\sigma^2_{\boldsymbol{Y}_i,\boldsymbol{X}_i,i}-\frac{1}{2}\mathbb{E}_{q(\boldsymbol{z}|\boldsymbol{x})}\left(\sum_{j=1}^{d}\frac{(Z_{ij}-\mu_{\boldsymbol{Y}_i,\boldsymbol{X}_i,j})^2}{\sigma^2_{\boldsymbol{Y}_i,\boldsymbol{X}_i,j}}\right)\\
=&-\frac{d}{2}\log(2\pi)-\frac{1}{2}\sum_{i=1}^{d}(\log\sigma^2_{\boldsymbol{Y}_i,\boldsymbol{X}_i,i}+1).
\end{aligned}
$$

**Jacobian regularization for stabilizing optimization.** To stabilize the training process and make the model robust against noises, we adopt the Jacobian regularization method from [13, 14]. Suppose $\boldsymbol{Z}_i \in \mathbb{R}^d$, $\boldsymbol{Y}_i \in \mathbb{R}^G$, $\boldsymbol{Z}_{ij}$ is the $j$-th coordinate of vector $\boldsymbol{Z}_i$ and $\boldsymbol{Y}_{ig}$ is the $g$-th coordinate of vector $\boldsymbol{Y}_i$. Then we control the Frobenius norm of the Jacobians of the output of the encoder $\boldsymbol{Z}_i$ with respect to the input $\boldsymbol{Y}_i$:

$$
\Omega_{\text{Jacobian}}(\boldsymbol{Y}_1,\cdots,\boldsymbol{Y}_N;\boldsymbol{X}_1,\cdots\boldsymbol{X}_N)=\sum_{i=1}^{N}\sum_{j=1}^{d}\sum_{g=1}^{G}\mathbb{E}_{q(\boldsymbol{Z}_i|\boldsymbol{Y}_i,\boldsymbol{X}_i)}\left[\left(\frac{\partial\boldsymbol{Z}_{ij}}{\partial\boldsymbol{Y}_{ig}}\right)^2\right]. \tag{E.15}
$$

**MMD regularization for removing conditional differences.** We can also include the maximum mean discrepancy (MMD) loss as utilized in SAUCIE [56] and scArches [57], which explicitly penalizes the distances between cells from any pairings we want to adjust in the latent space. Let $Z_1, Z_1', Z_2, Z_2'$ be independent samples from to two distributions $\mathcal{P}_1, \mathcal{P}_2$, then the MMD for these two distributions is:

$$
\mathcal{D}(\mathcal{P}_1,\mathcal{P}_2)=\mathbb{E}_{Z_1,Z_1'}[k(Z_1,Z_1')]+\mathbb{E}_{Z_2,Z_2'}[k(Z_2,Z_2')]-2\mathbb{E}_{Z_1,Z_2}[k(Z_1,Z_2)]
$$

where $k(\cdot,\cdot)$ is a kernel function. Empirically, if we want to integrate data from any two cell groups $\mathcal{P}_1$ and $\mathcal{P}_2$ which have $n_1$ and $n_2$ cells respectively, we punish large MMD in the latent space. The MMD loss for two groups can be defined as follows:

$$
L_{\text{MMD}}(\mathcal{P}_1,\mathcal{P}_2)=-\frac{1}{n_1^2}\sum_{i_1,i_2\in\mathcal{P}_1}k(\boldsymbol{Z}_{i_1},\boldsymbol{Z}_{i_2})-\frac{1}{n_2^2}\sum_{i_1,i_2\in\mathcal{P}_2}k(\boldsymbol{Z}_{i_1},\boldsymbol{Z}_{i_2})+\frac{2}{n_1n_2}\sum_{i_1\in\mathcal{P}_1,i_2\in\mathcal{P}_2}k(\boldsymbol{Z}_{i_1},\boldsymbol{Z}_{i_2})
$$

where $\boldsymbol{Z}_i$ is the latent vector for cell $i$. In practice, we employ a multi-scale RBF kernel, proposed in trVAE [38]. The multi-scale RBF kernel is defined as $k(\boldsymbol{Z}_{i_1},\boldsymbol{Z}_{i_2}) = \sum_{i=1}^{l} k(\boldsymbol{Z}_{i_1},\boldsymbol{Z}_{i_2},\lambda_i)$ where $k(\boldsymbol{Z}_{i_1},\boldsymbol{Z}_{i_2},\lambda_i) = \exp(-\lambda_i\|\boldsymbol{Z}_{i_1}-\boldsymbol{Z}_{i_2}\|^2)$ and $\lambda_i$ is a hyper-parameter. We set $l = 19$ by default and let $\{\lambda_i\}_{i=1}^{l} = [10^{-6}, 10^{-5}, 10^{-4}, 10^{-3}, 10^{-2}, 0.1, 1, 5, 10, 15, 20, 25, 30, 35, 100, 10^3, 10^4, 10^5, 10^6]$. Then, the MMD penalty for the whole dataset can be defined as:

$$
\Omega_{\text{MMD}}=\sum_{(s,t)\in\text{all pairs}}L_{\text{MMD}}(\mathcal{P}_s,\mathcal{P}_t).
$$

Suppose we wish to apply the MMD penalty to a set of $k$ cell groups (such as replicates) denoted by $\{\mathcal{P}_1,\mathcal{P}_2,\ldots,\mathcal{P}_k\}$. We can define a set of pairs as pairs $= \{(i,j) \mid 1 \le i < j \le k\}$ to represent all possible pairwise combinations of the $k$ groups.

##### S.2.2 Posterior Estimation

1. $\boldsymbol{Z}_i|\boldsymbol{Y}_i,\boldsymbol{X}_i$

   Since the approximate posterior distribution of $\boldsymbol{Z}_i|\boldsymbol{Y}_i,\boldsymbol{X}_i$ is $\mathcal{N}_d\left(\boldsymbol{\mu}_{\boldsymbol{Y}_i,\boldsymbol{X}_i},\boldsymbol{\Sigma}_{\boldsymbol{Y}_i,\boldsymbol{X}_i}\right)$, we can use $\boldsymbol{\mu}_{\boldsymbol{Y}_i,\boldsymbol{X}_i}$ as the latent representation of $\boldsymbol{Y}_i$.

2. $c_i|\boldsymbol{Y}_i,\boldsymbol{X}_i$

   It is worth noting that a useful property of our model is $\boldsymbol{Y}_i \perp\!\!\!\perp (w_i,c_i)|\boldsymbol{Z}_i,\boldsymbol{X}_i$, which is due to the fact that the probability mass function of $Y_{ig}|\boldsymbol{Z}_i,w_i,c_i,\boldsymbol{X}_i$ is $p(y|\boldsymbol{Z}_i,w_i,c_i,\boldsymbol{X}_i)=p(y|\boldsymbol{Z}_i,\boldsymbol{X}_i)$, which only depends on $\boldsymbol{Z}_i$ and $\boldsymbol{X}_i$. Note that we have already derived the formula for $p(\boldsymbol{z}|c_i=c)$ in Appendix S.2.1, we have

$$
\begin{aligned}
p(c|\boldsymbol{Y}_i,\boldsymbol{X}_i) &= \int_{\mathbb{R}} p(c,\boldsymbol{z}|\boldsymbol{Y}_i,\boldsymbol{X}_i)\mathrm{d}z \\
&\xlongequal{\boldsymbol{Y}_i|\boldsymbol{Z}_i,\boldsymbol{X}_i \perp\!\!\!\perp (w_i,c_i)|\boldsymbol{Z}_i,\boldsymbol{X}_i} \int_{\boldsymbol{z}} p(c|\boldsymbol{Z}_i=\boldsymbol{z})p(\boldsymbol{z}|\boldsymbol{Y}_i,\boldsymbol{X}_i)\mathrm{d}\boldsymbol{z} \\
&= \mathbb{E}_{p(\boldsymbol{Z}_i|\boldsymbol{Y}_i,\boldsymbol{X}_i)}\left[\frac{p(\boldsymbol{z}|c_i=c,\boldsymbol{X}_i)p(c,\boldsymbol{X}_i)}{p(\boldsymbol{z},\boldsymbol{X}_i)}\right] \\
&\xlongequal{(\boldsymbol{Z}_i,c_i) \perp\!\!\!\perp \boldsymbol{X}_i} \mathbb{E}_{p(\boldsymbol{Z}_i|\boldsymbol{Y}_i,\boldsymbol{X}_i)}\left[\frac{p(\boldsymbol{z}|c_i=c)p(c)}{p(\boldsymbol{z})}\right] \\
&\approx \frac{1}{L}\sum_{l=1}^{L}\frac{\widehat{p}(\boldsymbol{z}^{(l)}|c_i=c)\widehat{\pi}_c}{\sum\limits_{c'=1}^{K}\widehat{p}(\boldsymbol{z}^{(l)}|c_i=c')\widehat{\pi}_{c'}} \triangleq \widehat{p}(c|\boldsymbol{Y}_i,\boldsymbol{X}_i).
\end{aligned}
$$

   So we can obtain the predicted category of cell $i$ by computing the MAP estimator $\widehat{c}_i=\arg\max\limits_{c}\ \widehat{p}(c|\boldsymbol{Y}_i,\boldsymbol{X}_i)$.

3. $\tilde{\boldsymbol{w}}_i|\boldsymbol{Y}_i,\boldsymbol{X}_i$

   Note that

$$
p(w,c|\boldsymbol{Y}_i,\boldsymbol{X}_i) \approx \frac{1}{L}\sum_{l=1}^{L}\frac{\widehat{p}(\boldsymbol{z}^{(l)}|c_i=c,w_i=w)\widehat{\pi}_c}{\sum\limits_{c'=1}^{K}\widehat{p}(\boldsymbol{z}^{(l)}|c_i=c')\widehat{\pi}_{c'}} \triangleq \widehat{p}(w,c|\boldsymbol{Y}_i,\boldsymbol{X}_i).
$$

   Let $\boldsymbol{\mu}_{\tilde{\boldsymbol{w}}_i}$ and $\Sigma_{\tilde{\boldsymbol{w}}_i}$ denote the mean and the covariance matrix for the estimated distribution of $\tilde{\boldsymbol{w}}_i|\boldsymbol{Y}_i,\boldsymbol{X}_i$. We have

$$
\begin{aligned}
\boldsymbol{\mu}_{\tilde{\boldsymbol{w}}_i} &= \sum_{c=1}^{K}\int [wa_c+(1-w)b_c]\widehat{p}(w,c|\boldsymbol{Y}_i,\boldsymbol{X}_i)\mathrm{d}w \\
&\approx \tfrac{1}{M}\sum_{c=1}^{K}\sum_{j=1}^{M}\left[\tfrac{j}{M}a_c+\left(1-\tfrac{j}{M}\right)b_c\right]\tfrac{1}{L}\sum_{l=1}^{L}\frac{\widehat{p}(\boldsymbol{z}^{(l)}|c_i=c,w_i=\frac{j}{M})\widehat{\pi}_c}{\sum\limits_{c'=1}^{K}\widehat{p}(\boldsymbol{z}^{(l)}|c_i=c')\widehat{\pi}_c'}
\end{aligned}
$$

$$\mathrm{diag}(\Sigma_{\tilde{\boldsymbol{w}}_i}) \approx \frac{1}{M}\sum_{c=1}^{K}\sum_{j=1}^{M}\left[\frac{j}{M}a_c + \left(1-\frac{j}{M}\right)b_c\right]^2 \frac{1}{L}\sum_{l=1}^{L}\frac{\widehat{p}(\boldsymbol{z}^{(l)}|c_i = c, w = \frac{j}{M})\widehat{\pi}_c}{\sum_{c'=1}^{K}\widehat{p}(\boldsymbol{z}^{(l)}|c_i = c')\widehat{\pi}'_c} - \boldsymbol{\mu}^2_{\tilde{\boldsymbol{w}}_i}.$$

Here, as the integration over $w$ is intractable, we approximate the integral by the rectangular rule with equally spaced $M$ points in $[0, 1]$.

##### S.2.3 Edge Score

Below, seven kinds of edge scores are presented.

1. Edge Score Based on Posterior Mean of $c$.

   The first edge score is based on the posterior mean $\frac{1}{N}\sum_{i=1}^{N} p(c_i|\boldsymbol{Y}_i, \boldsymbol{X}_i)$. Let

$$\mathcal{C}_{j_1j_2} = \{i : \mathbb{P}(c_i \in \{C_{j_1j_1}, C_{j_1j_2}, C_{j_2j_2}\}|\boldsymbol{Y}_i, \boldsymbol{X}_i) > t_1\}$$

   where $t_1 = 0.5$ is the threshold parameter. We are considering $\mathcal{C}_{j_1j_2}$ instead of the full data set because not every cell contributes to the existence of the edge $e_{j_1j_2}$. Within the set $\mathcal{C}_{j_1j_2}$, proportion of cells at the edge $e_{j_1j_2}$ gives us one score

$$\mathbb{E}\left[\frac{1}{|\mathcal{C}_{j_1j_2}|}\sum_{i\in\mathcal{C}_{j_1j_2}} \mathbb{1}_{\{c_i = C_{j_1j_2}\}}\right] \approx \frac{1}{|\widehat{\mathcal{C}}_{j_1j_2}|}\sum_{i\in\widehat{\mathcal{C}}_{j_1j_2}} \frac{\widehat{p}\,(C_{j_1j_2}|\boldsymbol{Y}_i, \boldsymbol{X}_i)}{\widehat{p}(C_{j_1j_1}|\boldsymbol{Y}_i, \boldsymbol{X}_i) + \widehat{p}(C_{j_1j_2}|\boldsymbol{Y}_i, \boldsymbol{X}_i) + \widehat{p}(C_{j_2j_2}|\boldsymbol{Y}_i, \boldsymbol{X}_i)},$$

   which is denoted by $s^{\text{mean}}_{j_1j_2}$, where

$$\widehat{\mathcal{C}}_{j_1j_2} = \{i : \widehat{p}(C_{j_1j_1}|\boldsymbol{Y}_i, \boldsymbol{X}_i) + \widehat{p}(C_{j_1j_2}|\boldsymbol{Y}_i, \boldsymbol{X}_i) + \widehat{p}(C_{j_2j_2}|\boldsymbol{Y}_i, \boldsymbol{X}_i) > t_1\}.$$

2. Edge Score Based on Modified Posterior Mean of $c$.

   A modified version of $s^{\text{mean}}_{j_1j_2}$ is given by

$$s^{\text{modified\_mean}}_{j_1j_2} = \frac{\sum_{i\in\widehat{\mathcal{C}}_{j_1j_2}} \widehat{p}\,(C_{j_1j_2}|\boldsymbol{Y}_i, \boldsymbol{X}_i)}{\sum_{i\in\widehat{\mathcal{C}}_{j_1j_2}} [\widehat{p}(C_{j_1j_1}|\boldsymbol{Y}_i, \boldsymbol{X}_i) + \widehat{p}(C_{j_1j_2}|\boldsymbol{Y}_i, \boldsymbol{X}_i) + \widehat{p}(C_{j_2j_2}|\boldsymbol{Y}_i, \boldsymbol{X}_i)]}$$

   where an first-order approximation $\mathbb{E}\left(\frac{X}{Y}\right) \approx \frac{\mathbb{E}X}{\mathbb{E}Y}$ is used.

3. Edge Score Based on Raw MAP of $c$.

   Alternatively, we can also compute the edge score based on the MAP of $c_i$'s,

$$\tilde{c}_i = \underset{c\in\{1,2,\cdots,K\}}{\arg\max}\ \widehat{p}(c_i = c|\boldsymbol{Y}_i, \boldsymbol{X}_i).$$

   Then an edge score can be defined as the proportion of the number of cells at the edge $e_{j_1j_2}$ to the number of cells at the vertices $j_1, j_2$ or the edge $e_{j_1j_2}$:

$$s_{j_1j_2}^{\text{raw\_map}} = \frac{|\{i : \tilde{c}_i = C_{j_1j_2}\}|}{|\{i : \tilde{c}_i \in \{C_{j_1j_2, j_1 \neq j_2}\}\}|}.$$

4. Edge Score Based on MAP of $c$.

   We can also define the edge score of $c_i$ as the proportion of the number of cells at the edge $e_{j_1j_2}$ to the number of cells at the vertices $j_1, j_2$ or the edge $e_{j_1j_2}$:

$$s_{j_1j_2}^{\text{map}} = \frac{|\{i : \tilde{c}_i = C_{j_1j_2}\}|}{|\{i : \tilde{c}_i \in \{C_{j_1j_1}, C_{j_1j_2}, C_{j_2j_2}\}\}|}.$$

5. Edge Score Based on Modified MAP of $c$.

   Since the assigned categories by MAP of $c$ are generally at the edges instead of the nodes, the estimated denominator will be smaller than the real one, and $s_{j_1j_2}^{\text{map}}$ will be large when the number of cells in nodes $j_1$ and $j_2$ is small. To address this problem, we further modify the denominator to get the modified MAP edge score:

$$s_{j_1j_2}^{\text{modified\_map}} = \frac{|\{i : \tilde{c}_i = C_{j_1j_2}\}|}{|\{i : \boldsymbol{e}_{j_1}^{\top}\boldsymbol{\mu}_{\tilde{\boldsymbol{w}}_i} > 0.5 \text{ or } \boldsymbol{e}_{j_1}^{\top}\boldsymbol{\mu}_{\tilde{\boldsymbol{w}}_i} > 0.5\}|}.$$

6. Edge Score Based on MAP of $\mu_{\tilde{w}_i}$.

   Sometimes the large number of cell type $k$ will boost the dimension of $\widehat{p}(c_i = c|\boldsymbol{Y}_i, \boldsymbol{X}_i)$ to $\frac{k(k-1)}{2}$, which may cause the edge score inference sensitive to $\widehat{p}(c_i = c|\boldsymbol{Y}_i, \boldsymbol{X}_i)$ estimation. Therefore, in the dataset containing lots of cell types, we can compute the edge score from $\mu_{\tilde{w}_i}$, which only has $k$ dimension and also contains the position information. First, define a set $MAP(j_1, j_2)$ containing the cells that have the largest $\mu_{\tilde{w}_i}$ value in the vertices $j_1, j_2$ dimension.

$$MAP(j_1, j_2) := \{i : \arg\max_j e_j^T \mu_{\tilde{w}_i} = (j_1 \text{ or } j_2)\}$$

   Then, the edge score can be defined by the proportion of the number of cells that lie in the middle of two vertices to the number of cells that have the largest $\mu_{\tilde{w}_i}$ value in these vertices' dimensions:

$$s_{j_1,j_2}^{\text{w\_base}} = \frac{|\{i : abs(e_{j_1}^T\mu_{\tilde{w}_i} - e_{j_2}^T\mu_{\tilde{w}_i}) < 0.1, i \in MAP(j_1, j_2)\}|}{|MAP(j_1, j_2)|}$$

7. Edge Score Based on Modified MAP of $\mu_{\tilde{w}_i}$.

   When there exists some unbalanced cell type categories, $MAP(j_1, j_2)$ that only care about the 1st rank dimension of $\mu_{\tilde{w}_i}$ may cause the edge score unexpectedly small when one of the vertices has a large number of cells while the other only has a small number. Then we can modified $MAP(j_1, j_2)$ to consider the largest 2 dimension of $\mu_{\tilde{w}_i}$.

$$MAP2(j_1, j_2) := \{i : \underset{j' \subset [k], |j'|=2}{\arg\max} \sum_{j \in j'} e_j^T \mu_{\tilde{w}_i}, j' = \{j_1, j_2\}\}$$

Then, the edge score can be defined by the proportion of the number of cells that lie in the middle of the two vertices $j_1, j_2$ to the number of cells that belong to $MAP2(j_1, j_2)$:

$$s_{j_1,j_2} = \frac{|\{i : \max_j e_j^T \mu_{\tilde{w}_i} < 0.55 \cdot (e_{j_1} + e_{j_2})^T \mu_{\tilde{w}_i}), i \in MAP2(j_1, j_2)\}|}{|MAP2(j_1, j_2)|}$$

By default, the package uses edge scores based on the modified MAP of $c$.

#### S.2.4 Proof of Proposition 1

**Proof.** Note that

$$\mathbb{E}_{\widehat{p}(\tilde{\boldsymbol{w}}_i|\boldsymbol{Y}_i,\boldsymbol{X}_i)}\|\tilde{\boldsymbol{w}}_i - \boldsymbol{w}\|_2^2 = \mathbb{E}_{\widehat{p}(\tilde{\boldsymbol{w}}_i|\boldsymbol{Y}_i,\boldsymbol{X}_i)}(\tilde{\boldsymbol{w}}_i^\top \tilde{\boldsymbol{w}}_i) - 2\boldsymbol{w}^\top \tilde{\boldsymbol{w}}_i + \boldsymbol{w}^\top \boldsymbol{w}$$
$$\|\mathbb{E}_{\widehat{p}(\tilde{\boldsymbol{w}}_i|\boldsymbol{Y}_i,\boldsymbol{X}_i)}\tilde{\boldsymbol{w}}_i - \boldsymbol{w}\|_2^2 = (\mathbb{E}_{\widehat{p}(\tilde{\boldsymbol{w}}_i|\boldsymbol{Y}_i,\boldsymbol{X}_i)}\tilde{\boldsymbol{w}}_i)^\top \mathbb{E}_{\widehat{p}(\tilde{\boldsymbol{w}}_i|\boldsymbol{Y}_i,\boldsymbol{X}_i)}\tilde{\boldsymbol{w}}_i - 2\boldsymbol{w}^\top \tilde{\boldsymbol{w}}_i + \boldsymbol{w}^\top \boldsymbol{w},$$

we have

$$\begin{aligned}\mathbb{E}_{\widehat{p}(\tilde{\boldsymbol{w}}_i|\boldsymbol{Y}_i,\boldsymbol{X}_i)}\|\tilde{\boldsymbol{w}}_i - \boldsymbol{w}\|_2^2 = {} & \|\mathbb{E}_{\widehat{p}(\tilde{\boldsymbol{w}}_i|\boldsymbol{Y}_i,\boldsymbol{X}_i)}\tilde{\boldsymbol{w}}_i - \boldsymbol{w}\|_2^2 \\ & - (\mathbb{E}_{\widehat{p}(\tilde{\boldsymbol{w}}_i|\boldsymbol{Y}_i,\boldsymbol{X}_i)}\tilde{\boldsymbol{w}}_i)^\top \mathbb{E}_{\widehat{p}(\tilde{\boldsymbol{w}}_i|\boldsymbol{Y}_i,\boldsymbol{X}_i)}\tilde{\boldsymbol{w}}_i + \mathbb{E}_{\widehat{p}(\tilde{\boldsymbol{w}}_i|\boldsymbol{Y}_i,\boldsymbol{X}_i)}(\tilde{\boldsymbol{w}}_i^\top \tilde{\boldsymbol{w}}_i),\end{aligned}$$

where the last two terms do not involve $\boldsymbol{w}$. So, minimizing $\mathbb{E}_{\widehat{p}(\tilde{\boldsymbol{w}}_i|\boldsymbol{Y}_i,\boldsymbol{X}_i)}\|\tilde{\boldsymbol{w}}_i - \boldsymbol{w}\|_2^2$ is equivalent to minimizing $\|\mathbb{E}_{\widehat{p}(\tilde{\boldsymbol{w}}_i|\boldsymbol{Y}_i,\boldsymbol{X}_i)}\tilde{\boldsymbol{w}}_i - \boldsymbol{w}\|_2^2$ over $\boldsymbol{w}$. The optimization problem (10) is thus equivalent to the optimization problem (11).

We divide the problem into two cases: finding the best projection onto edges and the best projection onto vertices.

(1) For projection onto edges, the original problem becomes

$$\begin{aligned}\min_{(j_1,j_2)\in\mathcal{E}(\widehat{\mathcal{B}})} \quad & (\boldsymbol{w}_{j_1} - \mu_{j_1})^2 + (\boldsymbol{w}_{j_2} - \mu_{j_2})^2 - (\mu_{j_1}^2 + \mu_{j_2}^2) \\ \text{s.t.} \quad & \|\boldsymbol{w}\|_1 = w_{j_1} + w_{j_2} = 1, \quad \boldsymbol{w} \succeq \boldsymbol{0}_k.\end{aligned}$$

Notice that for any edge $(j_1, j_2) \in \mathcal{E}(\widehat{\mathcal{B}})$, the objective function is quadratic in $w_{j_1}$ (and $w_{j_2}$) since

$$\begin{aligned}&(w_{j_1} - \mu_{j_1})^2 + (w_{j_2} - \mu_{j_2})^2 - (\mu_{j_1}^2 + \mu_{j_2}^2) \\ =&(w_{j_1} - \mu_{j_1})^2 + (1 - w_{j_1} - \mu_{j_2})^2 - (\mu_{j_1}^2 + \mu_{j_2}^2) \\ =&2\left(w_{j_1} - \frac{1 + \mu_{j_1} - \mu_{j_2}}{2}\right)^2 + \frac{(1 - \mu_{j_1} - \mu_{j_2})^2}{2}\end{aligned}$$

$$\geq \frac{(1-\mu_{j_1}-\mu_{j_2})^2}{2},$$

with equality holds if and only if $w_{j_1} = \mu_{j_1} + \frac{1-\mu_{j_1}-\mu_{j_2}}{2}$. That is, for each edge $(j_1, j_2) \in \mathcal{E}(\widehat{\mathcal{B}})$, the solution

$$w_{k'} = \begin{cases} \mu_{j_1} + \frac{1-\mu_{j_1}-\mu_{j_2}}{2}, & k' = j_1 \\ \mu_{j_2} + \frac{1-\mu_{j_1}-\mu_{j_2}}{2}, & k' = j_2 \\ 0, & \text{otherwise,} \end{cases}$$

achieves minimum projection error

$$\|\boldsymbol{w} - \boldsymbol{\mu}_{\tilde{\boldsymbol{w}}_i}\|_2^2 = \sum_{k' \neq j_1, j_2} \mu_{k'}^2 + \frac{(1-\mu_{j_1}-\mu_{j_2})^2}{2}.$$

If we want edge $(j_1, j_2)$ to have the smallest projection error, then for any other edge $(m_1, m_2) \in \mathcal{E}(\widehat{\mathcal{B}})$, we need to have

$$\begin{aligned} & \|\text{proj}_{(j_1,j_2)}(\boldsymbol{\mu}_{\tilde{\boldsymbol{w}}}) - \boldsymbol{\mu}_{\tilde{\boldsymbol{w}}_i}\|_2^2 \leq \|\text{proj}_{(m_1,m_2)}(\boldsymbol{\mu}_{\tilde{\boldsymbol{w}}}) - \boldsymbol{\mu}_{\tilde{\boldsymbol{w}}_i}\|_2^2 \\ \iff \quad & \sum_{k' \neq j_1, j_2} \mu_{k'}^2 + \frac{(1-\mu_{j_1}-\mu_{j_2})^2}{2} \leq \sum_{k' \neq m_1, m_2} \mu_{k'}^2 + \frac{(1-\mu_{m_1}-\mu_{m_2})^2}{2} \\ \iff \quad & (\mu_{j_1}-\mu_{j_2})^2 + 2(\mu_{j_1}+\mu_{j_2}) \geq (\mu_{m_1}-\mu_{m_2})^2 + 2(\mu_{m_1}+\mu_{m_2}). \end{aligned} \tag{E.16}$$

Thus, the best-projected edge can be obtained by computing

$$(j_1^*, j_2^*) = \underset{(j_1,j_2)\in\mathcal{E}(\widehat{\mathcal{B}})}{\arg\max} \quad (\mu_{j_1}-\mu_{j_2})^2 + 2(\mu_{j_1}+\mu_{j_2}).$$

(2) For the vertex $j \in \mathcal{N}(\widehat{\mathcal{B}})$, the problem becomes

$$\min_{\boldsymbol{w}} \quad (w_j - \mu_j)^2 \qquad \text{s.t.} \quad \|\boldsymbol{w}\|_1 = w_j = 1, \;\; \boldsymbol{w} \succeq \boldsymbol{0}_k,$$

and the solution is $\text{proj}_j(\boldsymbol{\mu}_{\tilde{\boldsymbol{w}}}) = \boldsymbol{e}_j$. The projection error for the vertex $i$ is given by

$$\|\text{proj}_j(\boldsymbol{\mu}_{\tilde{\boldsymbol{w}}}) - \boldsymbol{\mu}_{\tilde{\boldsymbol{w}}}\|_2^2 = \sum_{k' \neq j} \mu_{k'}^2 + (\mu_j - 1)^2.$$

Then $j_3^* \in \underset{j\in\mathcal{N}(\widehat{\mathcal{B}})}{\arg\min} \|\text{proj}_j(\boldsymbol{\mu}_{\tilde{\boldsymbol{w}}}) - \boldsymbol{\mu}_{\tilde{\boldsymbol{w}}}\|_2^2$ gives the best projection onto vertices.

(3) Consider the following case,

$$\|\text{proj}_{j_3^*}(\boldsymbol{\mu}_{\tilde{\boldsymbol{w}}}) - \boldsymbol{\mu}_{\tilde{\boldsymbol{w}}_i}\|_2^2 \leq \|\text{proj}_{(j_1^*,j_2^*)}(\boldsymbol{\mu}_{\tilde{\boldsymbol{w}}}) - \boldsymbol{\mu}_{\tilde{\boldsymbol{w}}_i}\|_2^2$$

$$\iff \quad \sum_{k' \neq j_3^*} \mu_{k'}^2 + (1-\mu_{j_3^*})^2 \leq \sum_{k' \neq j_1^*, j_2^*} \mu_{k'}^2 + \frac{(1-\mu_{j_1^*}-\mu_{j_2^*})^2}{2}$$

$$\iff \quad \mu_{j_1^*}^2 + \mu_{j_2^*}^2 + (1-\mu_{j_3^*})^2 \leq \mu_{j_3^*}^2 + \frac{(1-\mu_{j_1^*}-\mu_{j_2^*})^2}{2},$$

$$\iff \quad 4\mu_{j_3^*} - 1 \geq (\mu_{j_1^*} - \mu_{j_2^*})^2 + 2(\mu_{j_1^*} + \mu_{j_2^*})$$

$$\iff \quad (\mu_{j_3^*} - \mu_{j_3^*})^2 + 2(\mu_{j_3^*} + \mu_{j_3^*}) - 1 \geq (\mu_{j_1^*} - \mu_{j_2^*})^2 + 2(\mu_{j_1^*} + \mu_{j_2^*}).$$

Define $\mathcal{EN}(\mathcal{G}) = \mathcal{E}(\mathcal{G}) \cup \{(j,j) : j \in \mathcal{N}(\mathcal{G})\}$, then

$$\underset{(j_1,j_2)\in\mathcal{EN}(\widehat{\mathcal{B}})}{\arg\max} \quad (\mu_{j_1} - \mu_{j_2})^2 + 2(\mu_{j_1} + \mu_{j_2}) - \mathbb{1}_{\{j_1=j_2\}}$$

gives solutions to both optimization problems. ■

#### S.3 Benchmarking

##### S.3.1 Benchmarking Datasets

Following the settings in [4], which provided a comprehensive overview and guideline for TI methods, we evaluate VITAE's performance in recovering six types of trajectory topologies, as shown in Fig. S.1. Our benchmarking datasets include 10 real scRNA-seq data and 15 synthetic datasets, summarized in Table 1.

The datasets from real scRNA-seq studies include 9 datasets from [4]. Among them, 5 datasets have "gold standard" labels according to [4] and are included to cover all trajectory types with at least 200 cells. As most "gold standard" datasets are small, we also include 4 extra datasets (*dentate*, *fibroblast*, *planaria_muscle*, *planaria_full*) with "silver standard" labels. The datasets and labels are all extracted from the Dyno platform [4], except for the dataset *dentate*, whose cells are mislabeled, and we directly extract the labels from the GEO database (accession number: GSE95315). For each dataset, the Dyno platform also provides its reference trajectory backbone and cell positions, from which we calculate reference pseudotime by our definition. In addition, to evaluate TI methods on a dataset with disconnected states, we create the dataset *immune* by combining purified 10085 B cells, 8385 CD56 NK cells, and 2612 CD14 Monocytes cells from [58].

The drawback of using real datasets for benchmarking is that only discrete cell labels are available, though the cells are experiencing continuous transitions. In other words, the true cell positions and ordering along the trajectory are only known at a low resolution. To better evaluate VITAE's performance in estimating the cell positions and pseudotime, we also include synthetic datasets for evaluation. We consider two different simulation approaches. One simulator we use is dyngen [18], a multi-modal simulation engine for studying dynamic cellular processes at single-cell resolution. dyngen is also used in [4] and provides a delicate way to generate scRNA-seq data starting from gene regulation and transcriptional factors. However, it is limited to generating only a few hundred genes. Thus, we only generate 1000 genes for each dyngen dataset and treat the dyngen datasets as non-UMI data as the generated counts are typically large.

We also generate four synthetic datasets from our own model framework. First, we train the model on a real data set to obtain a decoder and $\widehat{\boldsymbol{U}}$. Then, we generate each $\tilde{\boldsymbol{w}}_i$ and $\boldsymbol{Z}_i$ and the observed UMI counts $\boldsymbol{Y}_i$ following the hierarchical models ((E.3) and E.6). Specifically, we treat the estimated decoder and $\widehat{\boldsymbol{U}}$ as the true $f_g(\cdot)$ and $\boldsymbol{U}$ and design a trajectory backbone by connecting some edges between the vertices. In the four generated datasets, we do not include any confounding covariates $\boldsymbol{X}_i$. We use real dataset *dentate* to generate synthetic datasets *linear* and *bifurcation*, and the real dataset *fibroblast* to generate synthetic datasets *multifurcating* and *tree*.

##### S.3.2 Evaluation metrics

We use five different scores to measure TI methods' accuracy in recovering the true trajectory, cell positions, and pseudotime. All these scores range from 0 to 1, and a larger value indicates better performance.

First, to measure the difference between an estimated trajectory backbone $\widehat{\mathcal{B}}$ and the true trajectory backbone $\mathcal{B}$, we compute two scores: the GED score and the IM score, both of which evaluate the difference between two graphs and are invariant to the permutation of vertices. The graph edit distance [59] is defined as $\text{GED}(\widehat{\mathcal{B}}, \mathcal{B}) = \min_{e_1,\cdots,e_j \in \mathcal{P}(\widehat{\mathcal{B}},\mathcal{B})} \sum_{i=1}^{j} c(e_i)$ where $\mathcal{P}(\widehat{\mathcal{B}}, \mathcal{B})$ denotes the set of edit paths transforming graph $\widehat{\mathcal{B}}$ to the graph $\mathcal{B}$ and $c(e_i) \geq 0$ is the cost of an operation $e_i$. In other words, GED is a symmetric distance measure that quantifies the minimum number of graph edit operations needed to transform $\widehat{\mathcal{B}}$ into an isomorphic graph of $\mathcal{B}$. We then standardize the GED as

$$s_{\text{GED}} = 1 - \min\left\{\text{GED}(\widehat{\mathcal{B}}, \mathcal{B})/k, 1\right\},$$

so that it ranges between 0 and 1. Besides, following [4], we also compute the IM score based on the Ipsen-Mikhailov distance [60], $\text{IM}(\mathcal{B}, \widehat{\mathcal{B}})$, between the estimated and reference trajectory backbone. The IM distance is symmetric, measures the dissimilarity of adjacency matrices' spectra of the two graphs, and is bounded between 0 and 1. Our IM score is further defined as $s_{\text{IM}} = 1 - \text{IM}(\mathcal{B}, \widehat{\mathcal{B}})$.

Next, we measure the error of the estimated cell position $\widehat{\tilde{\boldsymbol{w}}}_i$ for each cell $i$. We use two scores, the adjusted rand index (ARI) and the generalized rand index (GRI), both of which are invariant under the permutation of vertices and allow an unequal number of vertices. The ARI [61] is a commonly used symmetric measure for the similarity between two clustering results. Following [4], the cells are assigned to their nearest discrete state based on their estimated $\widehat{\tilde{\boldsymbol{w}}}_i$ and true values of $\tilde{\boldsymbol{w}}_i$, and the two groupings are compared using ARI. Since such discretization only compares the accuracy of $\widehat{\tilde{\boldsymbol{w}}}_i$ at a low resolution, we also define a GRI score that directly compares $\widehat{\tilde{\boldsymbol{w}}}_i$ with $\tilde{\boldsymbol{w}}_i$, which is an extension of the rand index (RI). For any two pair of cells $i_1$ and $i_2$, we define its "true" similarity as $\rho_{i_1 i_2} = \langle \sqrt{\tilde{\boldsymbol{w}}_{i_1}}, \sqrt{\tilde{\boldsymbol{w}}_{i_2}} \rangle$ where $\sqrt{\boldsymbol{v}} = (\sqrt{v_1}, \cdots, \sqrt{v_k})$ for a vector $\boldsymbol{v}$ of length $k$. The square-root operation is to guarantee that $\rho_{i_1 i_2} = 1$ if and only if $\tilde{\boldsymbol{w}}_{i_1} = \tilde{\boldsymbol{w}}_{i_2}$. Similarly, we define the estimated similarity between the two cells as $\widehat{\rho}_{i_1 i_2} = \langle \sqrt{\widehat{\tilde{\boldsymbol{w}}}_{i_1}}, \sqrt{\widehat{\tilde{\boldsymbol{w}}}_{i_2}} \rangle$. Then to compare the similarities between the estimated and reference cell positions on the trajectory backbone, the GRI is defined as

$$\text{GRI} = \sum_{i_1=1}^{n-1} \sum_{i_2=i_1+1}^{n} (1 - |\rho_{i_1 i_2} - \widehat{\rho}_{i_1 i_2}|) \Big/ \binom{2}{n}.$$

The rand index is a special case of GRI when all cells are positioned exactly on vertices.

Finally, we also measure the similarity between the reference and estimated pseudotime of cells along the trajectory, which is simply the Pearson correlation between the estimated pseudotime $\widehat{T}_i$ and the reference pseudotime $T_i$ across cells. Specifically, we define the PDT score ranging between 0 and 1 as $s_{\text{PDT}} = \left(\text{Cor}(\boldsymbol{T}, \widehat{\boldsymbol{T}}) + 1\right)/2$.

##### S.3.3 DE analysis

We compare VITAE with tradeSeq [16] and PseudotimeDE [17] on differentially expressed testing. Similar to [16], we use package dyntoy from dynverse toolbox [18] to generate linear trajectories. We perform 10 simulations. For each simulation, we generate a synthetic scRNA-seq dataset with 5000 genes and varying numbers of cells in $n \in \{250, 500, 750, 1000\}$. We set the differentially expressed rate to be 0.2, so that

20% of the genes are active. Then, we apply both Slingshot and VITAE on the simulated datasets to infer trajectory and estimate the pseudotime for all cells. We next apply DE test methods VITAE, tradeSeq, and PseudotimeDE on the estimated pseudotime with BH procedure and significance level 5%. We provide the details of the configuration below.

For Slingshot, we follow the usual preprocessing procedure to use all the genes for dimension reduction with PCA and clustering the cells with the default parameters of Seurat. Then we extract the principle components, cluster assignment for all cells, and the initial cluster as the input to Slingshot. For tradeSeq, we follow their workflow to call `fitGAM` function with `nknots=4` and use `associationTest` function with `contrastType='consecutive'`, which returns the adjusted $p$-values for all genes. For PseudotimeDE, we use 50 random subsampled raw count datasets of size $0.8n$, and apply Singshot to estimate the pseudotime for each of the datasets. Then, we call `runPseudotimeDE` function with parameter `model = 'nb'` and 10 CPU cores to obtain the adjusted $p$-values. We evaluate the three DE test methods using pseudotime estimated from trajectory inference methods Slingshot and VITAE. We did not explore PseudotimeDE based on VITAE's estimated pseudotime due to the computational cost. Therefore, our comparison consists of five combinations of TI and DE methods: Slingshot+VITAE, Slingshot+tradeSeq, Slingshot+PseudotimeDE, VITAE+VITAE, and VITAE+tradeSeq.

The results of the false discovery rate (FDR) and true positive rate (TPR) are shown in Fig. S.2. The signals are strong in the synthetic data (the signal strength is not adjustable in dyntoy), so all methods have powers of almost 1. In terms of FDR control, we have better control than tradeSeq. Compared with PseudotimeDE, our method has similar FDR control but is less computationally expensive. In terms of computational complexity, for a dataset with 1000 cells and 5000 genes, it took about 1 min, 2 min, and 2 h for VITAE+VITAE, Slingshot+tradeSeq, and Slingshot+PseudotimeDE to perform both trajectory inference and testing.

### S.4 Real data

#### S.4.1 Experiments on mouse brain datasets

To adjust for the confounding batch effect, we conducted 10 random trials on the joint analysis of two mouse neocortex datasets (Yuzwa and Ruan datasets), both with ($\gamma = 1$) and without ($\gamma = 0$) this loss, keeping all other hyperparameters at their default settings. As a stability measure of the estimated cell state centers in the latent space (denoted as $\boldsymbol{U}$) and the estimated probabilities for a cell to choose a vertex/edge (denoted as $\boldsymbol{\pi}$), we calculated the standard deviation of the pairwise distances between columns of $\boldsymbol{U}$ (pairwise distances between cluster centers) and the standard deviation of each entry in $\boldsymbol{\pi}$ across the 10 trials. In Fig. S.3, we observe a substantial improvement in the stability of our estimates with the Jacobian regularizer.

##### S.4.2 Experiments on the integration of three mouse brain datasets

**Experiment setup.** To adjust for confounding batch effect, we include the dataset ID of each cell (whether it is from Di Bella's, Yuzwa's, or Ruan's dataset) as covariates. In addition, as the Apical Progenitors are experiencing cell division, we also adjust for the cell-cycle effects by adding the cell-cycle scores (`S.Score` and `G2M.Score`) provided by their own datasets. To perform the joint trajectory analysis, we apply VITAE_Gauss with the aforementioned covariates. We provide the annotated cell types to initialize the latent space after the pretraining step. For the model hyperparameters, we use 96 PCs, two hidden layers with units 48 and 24 for the encoder, a symmetric structure for the decoder, and 16 units for the latent space. Since the dataset is larger and more complicated, we retain more PCs as input and use a wider network structure. For pre-training and training, we set $\gamma = 1$ for the Jacobian regularizers.

**Integration.** We further evaluated the integration capabilities of VIATE against other baseline methods, including Seurat CCA [24], Monocle 3 [15], scVI [26], and Scanorama [27]. The latter two are among the leading integration methods that are suitable for downstream trajectory inference [62]. The VIATE model setup is detailed in the preceding section. For Seurat CCA, we employed the `Seurat V3` R package and the `RunCCA` function. With `Monocle 3`, the `align_cds` function was utilized. For scVI, source information was incorporated using the `batch_key` parameter in the `scvi.model.SCVI` function. Lastly, Scanorama integration was achieved through the `scanorama.correct_scanpy` function. All integration processes are succeeded by the application of UMAP to derive a 2-dimensional embedding of the integrated dataset.

**Evaluation metric for integration.** We construct a mixing score based on the reduced UMAP space to measure the mixing effect of Day E18 Replicate, Day P1 Replicate, different days, and different cell types within the same day. We first calculate the KNN graph in UMAP space. Suppose we have $n$ categories, and we want to quantify how these $n$ categories are mixed. In the $k$ neighbors of a cell, assume there are $x$ cells that have the same category as this cell; then, for each cell, the mixing metric can be calculated by

$$\text{mixing score} = 1 - \frac{(x - \frac{k}{n})^+ + [k - \frac{k}{n} - (k - x)]^+}{2(1 - \frac{1}{n})k},$$

where $(x - a)^+ = \max\{x - a, 0\}$. The score reaches 1 when $x \le k/n$ and reaches 0 when $x = k$. The denominator $2(1 - n^{-1})k$ is used to normalize the score between 0 and 1. This score can also be converted to quantify the separation of different categories:

$$\text{separation score} = 1 - \text{mixing score}.$$

Consequently, we use the mixing score to measure the mixing effect of Day E18 Replicate, Day P1 Replicate, and different days. The separation score is applied to quantify the biological cell type conservation on different days.

##### S.4.3 Experiments on multi-omic human hematopoiesis datasets

To adjust for confounding batch effect, we include the sequencing technique of each cell (scRNA-seq or scATAC-seq) and also use MMD loss calculated within each annotated cell type to remove effects from different sequencing techniques better. For the model hyperparameters, we use 128 PCs, two hidden layers with units 32 and 16 for the encoder, a symmetric structure for the decoder, and 8 units for the latent space. For pre-training, we set $\gamma = \kappa = 0.6$ for the Jacobian and MMD regularizers. For training, we set $\gamma = \kappa = 1$ for the Jacobian and MMD regularizers.

We compare the trajectory results of VITAE to VIA [32] v0.1.96. Following the procedure in the original VIA paper, which involves employing Seurat CCA for cross-modality data integration and subsequently applying VIA to the merged dataset for trajectory analysis, we compare the visualization, inferred trajectory structure and pseudotime results between VIA and VITAE. As shown in Fig. S.8c, the inferred trajectory of VIA is less interpretable compared to VITAE. This is partially contributed by the Seurat's CCA integration method, which tends to overcorrect the cell embeddings, particularly affecting cell types with a limited number of cells, such as in the B cell lineage. Additionally, VIA fails to provide meaningful separations between some cell types even though the annotated cell types are provided as the input. Moreover, even when two cell types are visually distinct to each other, such as Baso.Eryth and Pre.B, VIA still incorrectly infers a connection between them.

#### S.5 Analysis of hyperparameters sensitivity and computational efficiency

##### S.5.1 Hyperparameter $\gamma$ for Jacobian regularization

We conducted 10 random trials on the joint analysis of the two mouse neocortex datasets (datasets from Yuzwa et. al [22] and Ruan et. al [23]), both with ($\gamma = 1$) and without ($\gamma = 0$) this loss, keeping all other hyperparameters at their default settings. As a stability measure of the estimated cell state centers ($\boldsymbol{U}$) in the latent space, we calculated the standard deviation of the pairwise distances between columns of $\boldsymbol{U}$ (pairwise distances between cluster centers) across the 10 trials. We also measure the stability of the estimated prior probabilities for a cell to choose each vertex/edge ($\boldsymbol{\pi}$), by calculating the standard deviation of each entry of $\boldsymbol{\pi}$ across the 10 trials. In Fig. S.3, we observe a substantial improvement in the stability of our estimates with the Jacobian regularizer.

##### S.5.2 Hyperparameter $\alpha$ for the Soft penalty term

We vary the value of the hyperparameter $\alpha$ on the soft penalty term in the joint analysis of the two mouse neocortex datasets (datasets from Yuzwa et. al [22] and Ruan et. al [23]), keeping all other hyperparameters in their default values. Fig. S0.2 shows the embedding of cells and the inferred trajectory after training with different $\alpha$ values: $\alpha = 0, 0.1, 0.15, 0.2, 0.5, 1$. We observe that, with the soft penalty terms, VITAE is more accurate in learning the shared trajectory of the two datasets. Its performance is relatively robust to minor adjustment of $\alpha$.

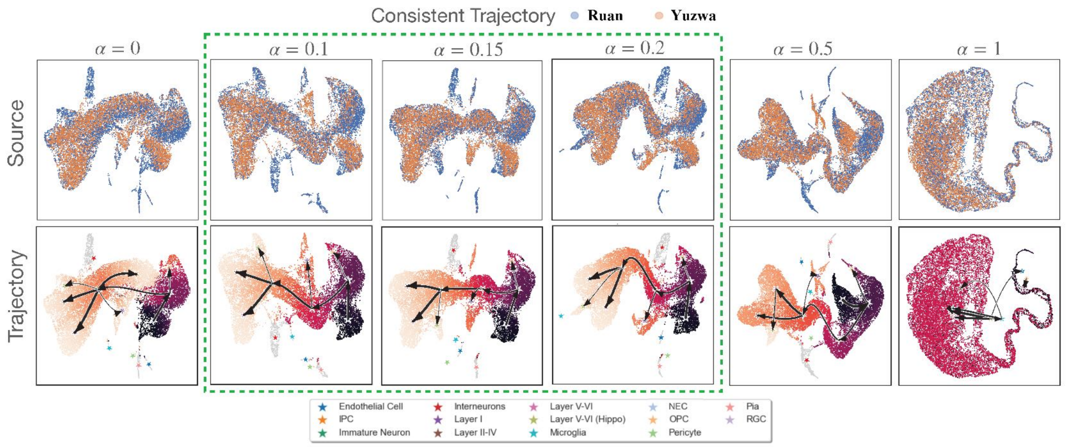

**Fig S0.2:** Sensitivity of VITAE on parameter $\alpha$ of soft penalty term.

##### S.5.3 Hyperparameter $\kappa$ for MMD regularization

We investigate the effect of varying the MMD loss penalty $\kappa$ in the joint trajectory analysis of scATAC-seq and scRNA-seq hematopoiesis data where the MMD loss is used to encourage the merging of two datasets. Fig. S0.3 shows the embedding of cells and the inferred trajectory after training with $\kappa$ values ranging from 0 to 1.5, keeping all other hyperparameters unchanged. Without MMD loss, the cells from the scRNA-seq and the scATAC-seq datasets cannot merge well. On the other hand, if $\kappa$ is too large, even though the cells can merge well, the latent representations are less dispersed, making it hard to infer the trajectory correctly. VITAE is relatively insensitive to $\kappa$ if $\kappa$ varies within a certain range.

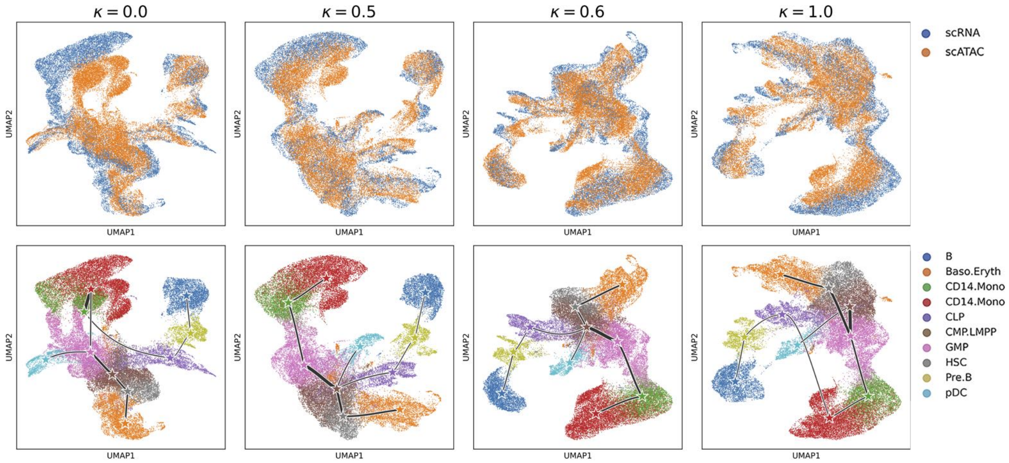

**Fig S0.3:** Sensitivity of VITAE on parameter $\kappa$ of MMD loss.

#### S.5.4 Computation efficiency and memory usage

We compare the computational efficiency of VITAE and alternative approaches when performing joint trajectory analysis on multiple datasets. Because Slingshot takes the dimension-reduced object as the input, it relies on data integration methods, such as Seurat's CCA, to adjust for potential batch effects. Therefore, it is more reasonable to compare the time complexity of both integration and trajectory inference.

To this end, we compared four different approaches:

1. VITAE that adjusts for both the data source differences and cell cycle scores;
2. Monocle with data source correction but no cell cycle score adjustments (as it can not adjust for continuous cell cycle scores);
3. Seurat that adjusts for both the data source differences and cell cycle scores to integrate the two datasets;
4. Seurat with data source correction but no cell cycle score adjustments.

The last two methods above provide an estimate of the shortest time for running Slingshot on the data integrated by Seurat. To evaluate these methods on different sizes of data, we subsampled cells from the joint datasets from Yuzwa et. al [22] and Ruan et. al [23], and generated a sequence of datasets of increasing cell numbers $n = 1000, 2000, \ldots, 15000, 16651$. As shown in Fig. S0.4, though VITAE is slightly slower than Monocle, it is faster than Seurat when adjusting for both batch effects and cell cycle scores.

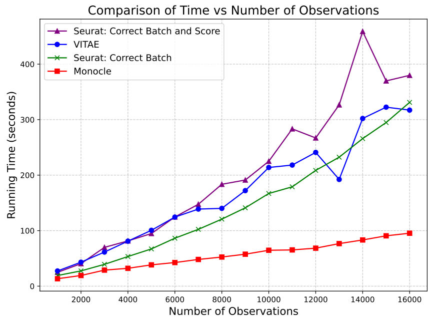

**Fig S0.4:** Computational time in the size of datasets for different methods.

We also record the memory usage of VITAE in real data analysis. The joint analysis of the two mouse neocortex data (Yuzwa et. al [22] and Ruan et. al [23]) is successfully executed on an 11th Gen Intel Core i7 CPU with 16 GB of RAM. The joint trajectory analysis of three datasets from Di Bella et. al [25], Yuzwa et. al [22], and Ruan et. al [23] can be performed on an Intel Xeon Gold CPU with 32 GB of RAM and a total running time of approximately 27 minutes.

#### S.6 Supplementary tables

**Table 1:** Detailed summary of datasets used in experiments, including trajectory topology, number of cells $N$, number of genes $G$, number of cell types $k$, and sources of the dataset.

| Type | Name | Count Type | Topology | $N$ | $G$ | $k$ | Source |
|---|---|---|---|---|---|---|---|
| real | aging | non-UMI | linear | 873 | 2815 | 3 | [63] |
| | human_embryos | non-UMI | linear | 1289 | 8772 | 5 | [64] |
| | germline | non-UMI | bifurcation | 272 | 8772 | 7 | [65] |
| | mesoderm | non-UMI | tree | 504 | 8772 | 9 | [66] |
| | cell_cycle | non-UMI | cycle | 264 | 6812 | 3 | [64] |
| | dentate | UMI | linear | 3585 | 2182 | 5 | [67] |
| | fibroblast | non-UMI | bifurcation | 355 | 3379 | 7 | [68] |
| | planaria_muscle | UMI | bifurcation | 2338 | 4210 | 3 | [7] |
| | planaria_full | UMI | tree | 18837 | 4210 | 33 | [7] |
| | immune | UMI | disconnected | 21082 | 18750 | 3 | [58] |
| synthetic | linear_1 | non-UMI | linear | 2000 | 991 | 4 | dyngen |
| | linear_2 | non-UMI | linear | 2000 | 999 | 4 | dyngen |
| | linear_3 | non-UMI | linear | 2000 | 1000 | 4 | dyngen |
| | bifurcating_1 | non-UMI | bifurcation | 2000 | 997 | 7 | dyngen |
| | bifurcating_2 | non-UMI | bifurcation | 2000 | 991 | 7 | dyngen |
| | bifurcating_3 | non-UMI | bifurcation | 2000 | 1000 | 7 | dyngen |
| | trifurcating_1 | non-UMI | multifurcating | 2000 | 969 | 10 | dyngen |
| | trifurcating_2 | non-UMI | multifurcating | 2000 | 995 | 10 | dyngen |
| | converging_1 | non-UMI | bifurcation | 2000 | 998 | 6 | dyngen |
| | cycle_1 | non-UMI | cycle | 2000 | 1000 | 3 | dyngen |
| | cycle_2 | non-UMI | cycle | 2000 | 999 | 3 | dyngen |
| | cycle_3 | non-UMI | cycle | 2000 | 999 | 3 | dyngen |
| | linear | UMI | linear | 1900 | 1990 | 5 | our model |
| | bifurcation | UMI | bifurcation | 2000 | 606 | 6 | our model |
| | multifurcating | UMI | multifurcating | 2000 | 606 | 6 | our model |
| | tree | UMI | tree | 2000 | 606 | 6 | our model |

**Table 2:** Number of cells on different embryonic days.

| Embryonic Day | E10.5 | E11.5 | E12.5 | E13.5 | E14.5 | E15.5 | E16.5 | E17.5 | E18.5 |
|---|---|---|---|---|---|---|---|---|---|
| Ruan's Dataset | 1172 | 0 | 2668 | 0 | 3742 | 1014 | 387 | 0 | 1278 |
| Yuzwa's Dataset | 0 | 1418 | 0 | 1137 | 0 | 2955 | 0 | 880 | 0 |

#### S.7 Supplementary figures

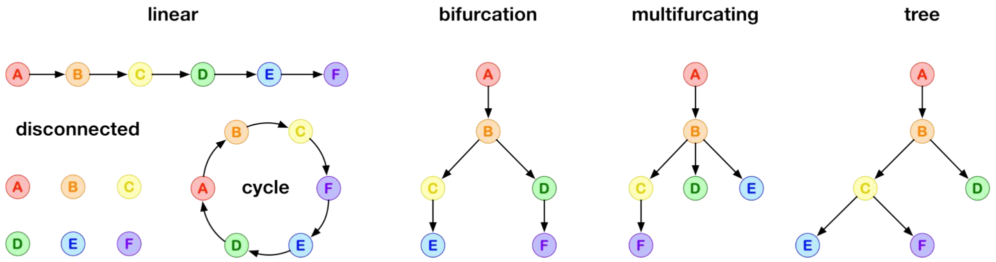

**Figure S.1:** Six different topologies of the underlying trajectories in real and synthetic datasets.

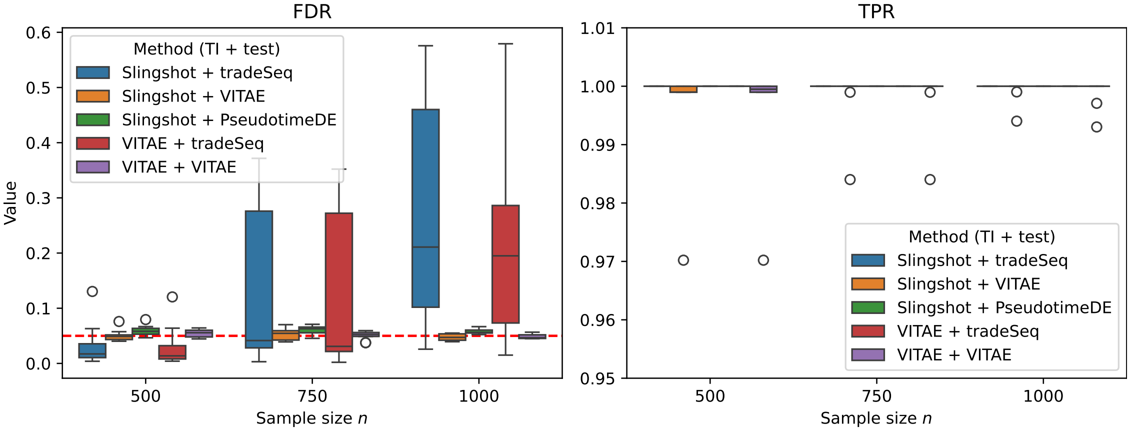

**Figure S.2:** Differential expressed testing results on simulated datasets generated by dyntoy. The false discovery ratio (FDR) and true positive ratio (TPR) are shown in the left and right panels, respectively.

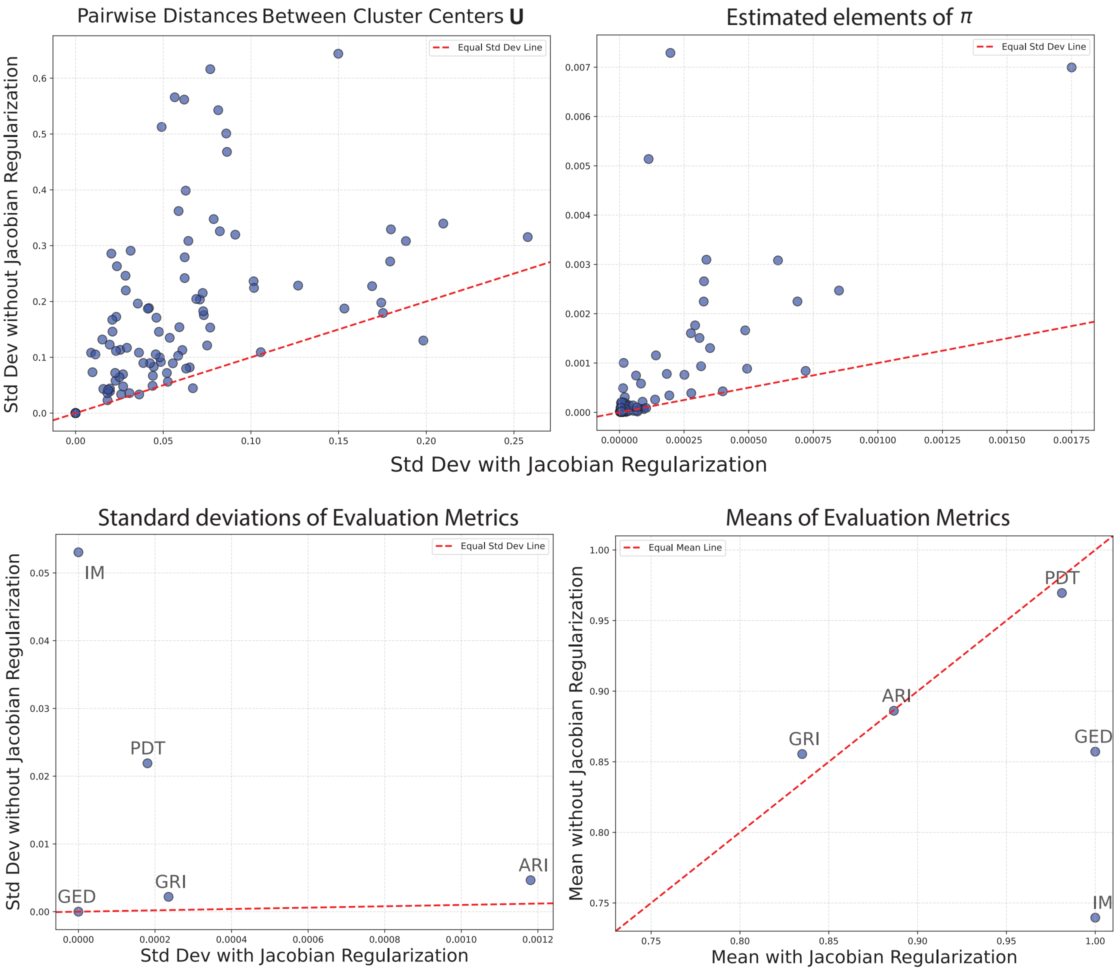

**Figure S.3: Comparison of stability with/without the Jacobian regularization.** VITAE is applied to perform a joint analysis of Yuzwa and Ruan datasets with 10 repeated trials with random initialization. Top left: standard derivations across 10 trials for the pairwise distances of the estimated cluster centers $U$ in the latent space. Top right: standard derivations across 10 trials for the estimated probabilities being assigned to each edge and vertex ($\pi$). Bottom left: standard derivations across 10 trials for the evaluation metric values between each trial's final result and the illustrated trajectory inference result in Figure 3. Bottom right: mean across 10 trials for the evaluation metric values between each trial's final result and the illustrated trajectory inference result in Figure 3.

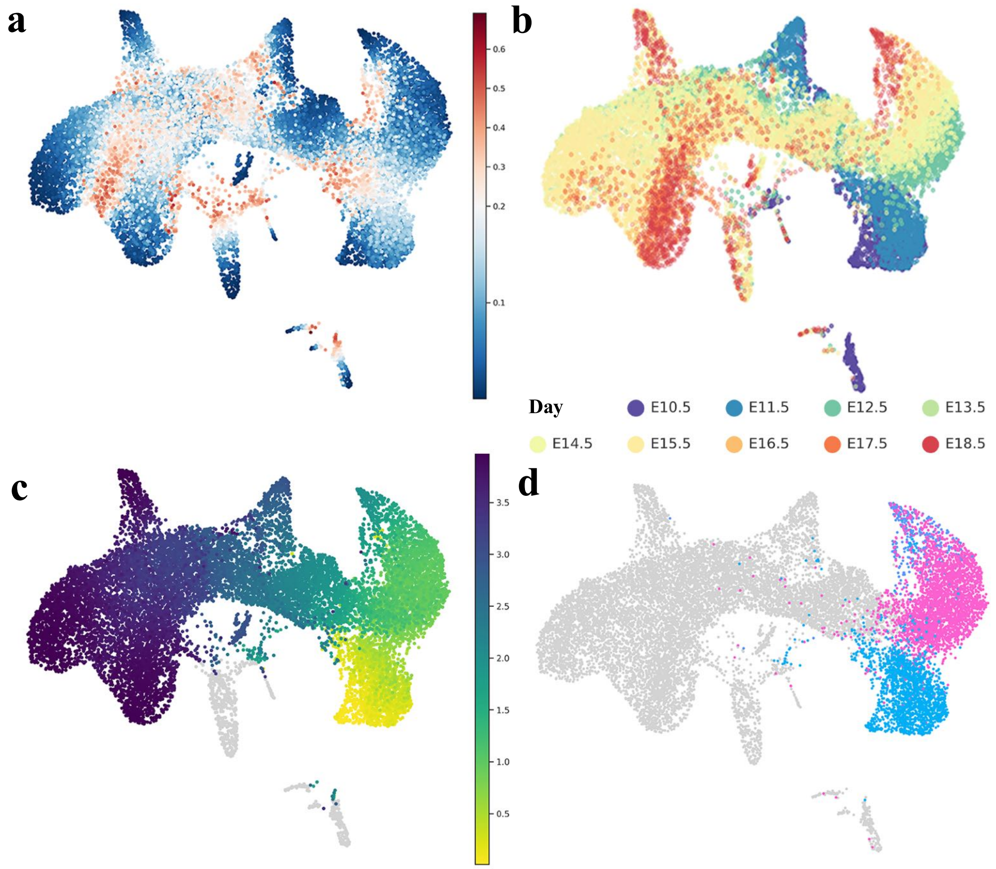

**Figure S.4: UMAP visualizations of VITAE's low-dimensional cell representations on the joint analysis of Yuzwa and Ruan datasets.** **(a)** Low-dimensional embeddings colored by the projection uncertainty estimated by VITAE. When projected to the inferred trajectory, Cells colored in red have higher uncertainties compared to cells colored in blue. **(b)** Low-dimensional embeddings colored by collection days. **(c)** Low-dimensional embeddings colored by the pseudotime estimated by VITAE. Cells not in the inferred trajectory are colored in gray. **(d)** Low-dimensional embeddings highlighting the NEC-RGC-OPC sub-trajectory where the cell types NEC, RGC, and OPC are highlighted.

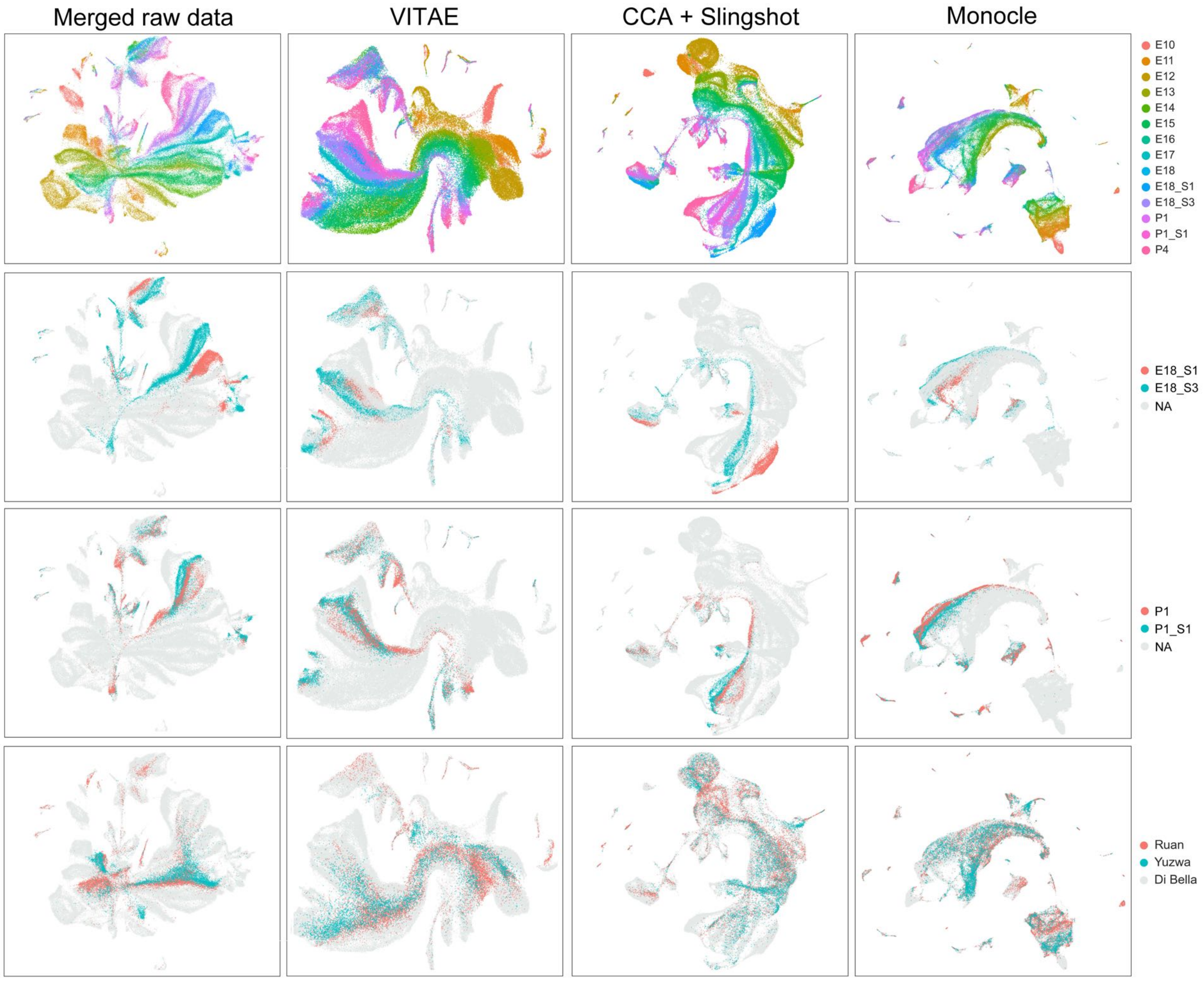

**Figure S.5: UMAP visualizations of the joint analysis of Di Bella, Yuzwa and Ruan datasets.** Each column represents a method. For the first column, cells are normalized and then concatenated to perform a joint PCA analysis before UMAP. In the first row and last row, cells are colored by their collection days and sources. In the second and third rows, cells from the two replicates at E18 and two replicates at P1 from the Di Bella dataset are highlighted.

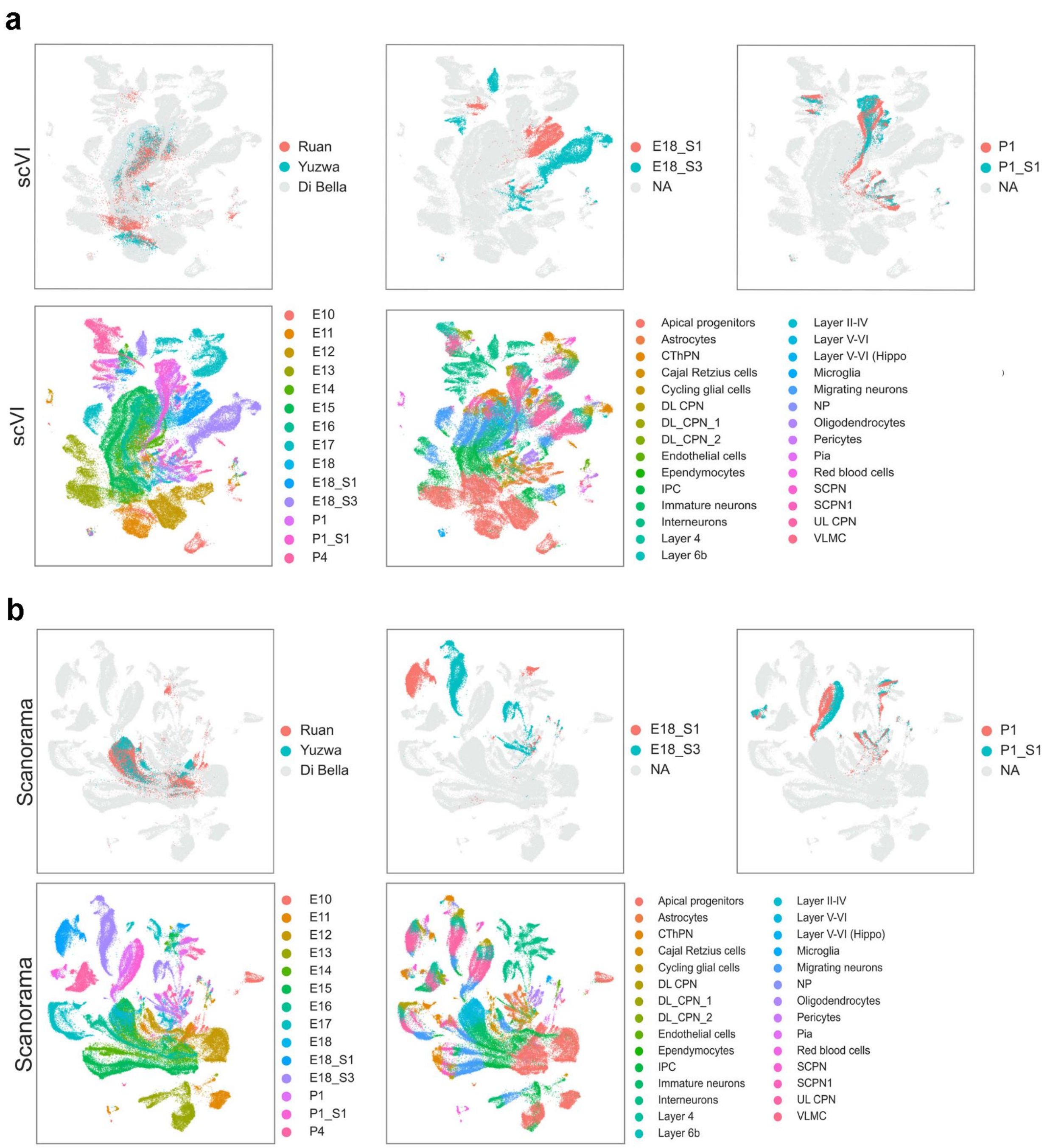

**Figure S.6: UMAP visualizations of the integration of Di Bella, Yuzwa and Ruan datasets using scVI and Scanorama.**

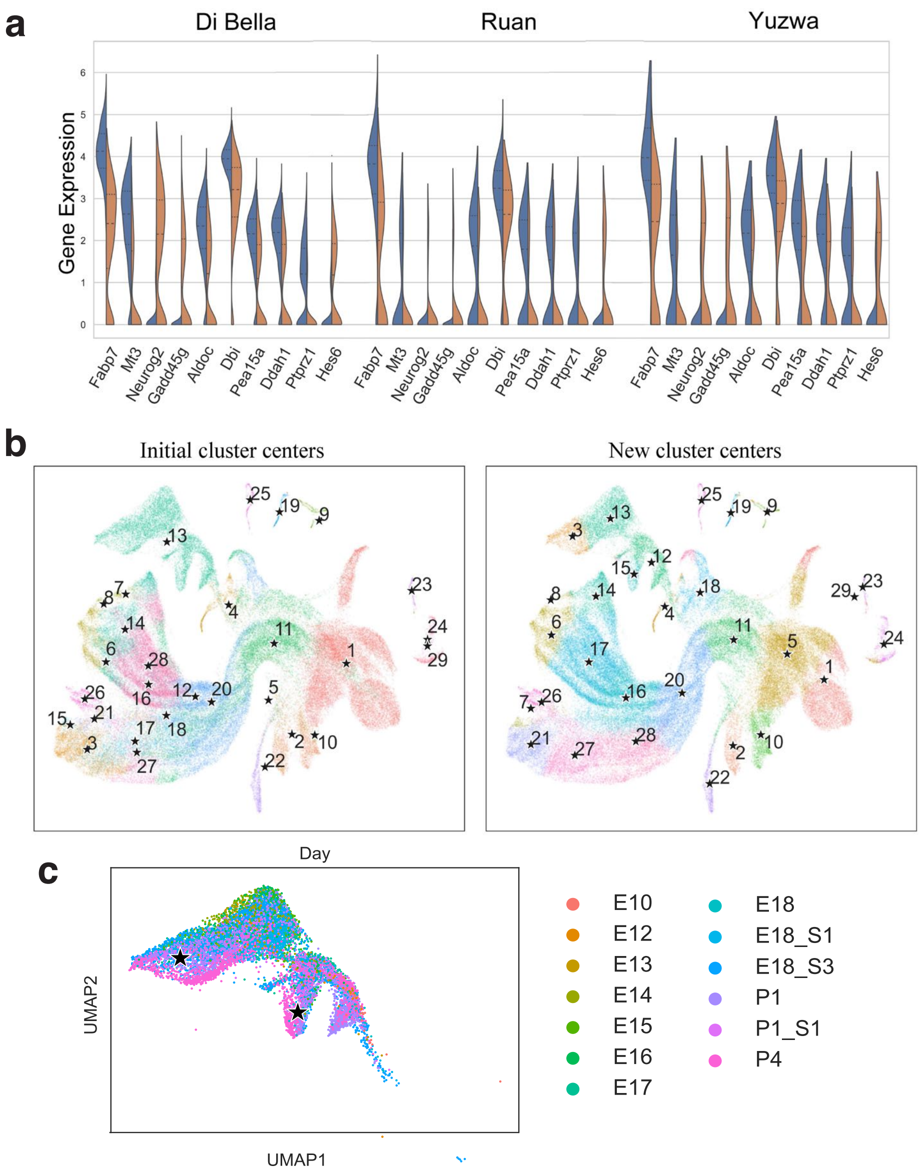

**Figure S.7: Additional results for joint trajectory analysis of Di Bella, Yuzwa and Ruan datasets. (a)** The violin plots of the top 10 differentially expressed gene expressions between the APs in the glial and neuronal branches in Fig. 4c. Tests are done by combining all three datasets while gene expressions are visualized for each dataset separately. **(b)** Locations of the cluster centers before and after VITAE training for the joint analysis of Di Bella, Yuzwa, and Ruan datasets. VITAE uses the reference cell types to initialize the locations of the vertices before training, and then the vertices will change locations and give new clustering results after training. The left and the right panels show the initial and trained cluster centers, respectively. For interneurons, though only one vertex (cluster 13) is initialized, VITAE can automatically learn four cluster centers after training. **(c)** Interneurons colored by collection days.

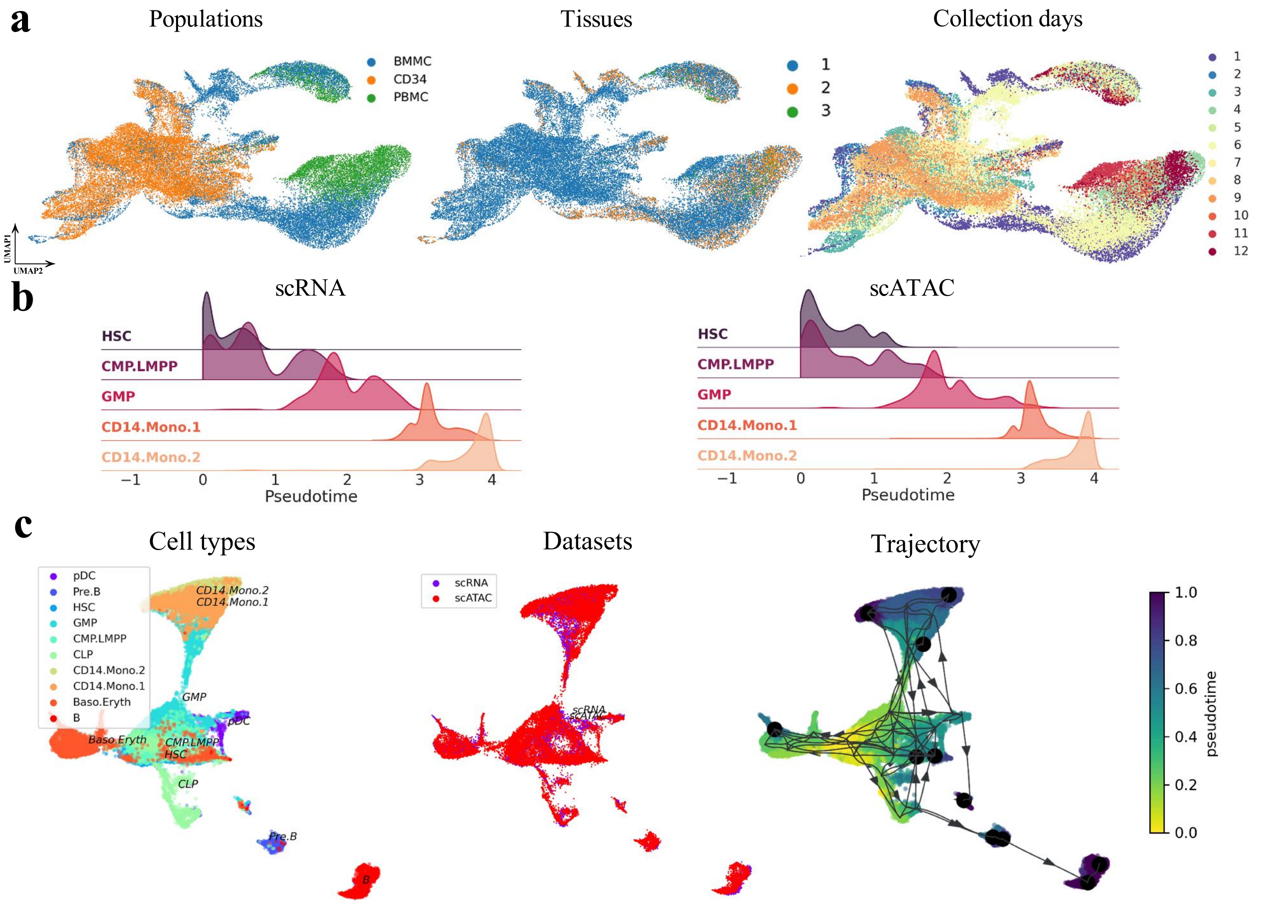

**Figure S.8: Integrative trajectory inference of multi-omic human hematopoiesis data.** **(a)** UMAP visualization of VITAE's low-dimensional embedding of cells, colored by cell populations ('BMMC', 'CD34', and 'PBMC' stand for bone marrow mononuclear cells, $CD34^{+}$-enriched bone marrow mononuclear cells, and peripheral blood mononuclear cells, respectively), tissues, replicates, and collection days. VITAE merges the gene expressions (scRNA) and the gene activities (scATAC) score, retains meaningful biological variations and correctly infers the developmental trajectory. **(b)** Distribution of pseudotime of the monocytic lineage for scRNA-seq and scATAC-seq cells. **(c)** Trajectory inference results of VIA [32].
